# Strength in Unity: a Dual Strategy to Restore NK Cell Cytotoxicity against Pancreatic Ductal Adenocarcinoma

**DOI:** 10.64898/2026.02.09.704789

**Authors:** Camille Rolin, Aubin Pitiot, Gilles Iserentant, Anaïs Oudin, Jean-Yves Servais, Victoria El-Khoury, Vanessa Barthelemy, Céline Hoffmann, Anna Golebiewska, Yong-Jun Kwon, Jacques Zimmer, Carole Seguin-Devaux

## Abstract

**Background:** Pancreatic ductal adenocarcinoma (PDAC), a condition representing 90% of pancreatic cancers, shows one of the lowest 5-year survival rates across all cancer types. Current therapeutic approaches remain largely inefficient, in part due to the presence of a hostile tumor microenvironment (TME), impeding immune cells infiltration and function. Specifically, Natural Killer (NK) cells from PDAC patients exhibit impaired phenotype and cytotoxic functions. NK cell immunotherapy represents a safe and promising approach to restore NK cell cytotoxicity against PDAC.

**Methods:** We developed a dual strategy based on i) the re-activation of NK cells through Natural Killer activating multimeric immunotherapeutic complexes (NaMiX) composed of IL-15/IL-15Rα dimers coupled to anti-NKp46 single-chain variable fragments (scFvs) and ii) the crosslinking of activated NK cells to PDAC cells with a Trispecific Killer Engager (TriKE) targeting NKG2D, NKp30 and the tumor-associated antigen CEA. We evaluated the ability of these constructs to stimulate NK cell functions across BxPC-3 PDAC cell line and patient-derived organoid models and in humanized NSG mice bearing PDAC xenografts.

**Results:** NaMiX stimulated the activation and cytotoxic functions of NK cells towards pancreatic BxPC-3 cells in vitro while TriKE cross-linked NK cells to BxPC-3 cells. The cytotoxic effects of NaMiX were further enhanced when combined with the crosslinking abilities of TriKE for the killing of NK cell-mediated BxPC-3 spheroid and PDAC patient-derived organoids. In humanized mice bearing BxPC-3 xenografts, NaMiX induced cytotoxic lymphocyte expansion, and increased tumor infiltration of NK cells, while TriKE tended to slow tumor progression.

**Conclusions:** This proof-of-concept study reports for the first time that activating and engaging NK cells with immunoconjugates are a promising therapeutic avenue for PDAC treatment. Efforts should now focus on the optimization of NK cell therapeutic modalities to favor the infiltration of a high number of NK cells into the tumor.

## BACKGROUND

Pancreatic ductal adenocarcinoma (PDAC) accounts for around 90% of pancreatic cancer cases and represents one of the deadliest cancer types.^1^ Delayed diagnosis as well as the extremely hostile tumor microenvironment (TME) render current therapeutic approaches inefficient and lead to a dramatic 5-year survival rate of only 13.3%.^2^ In particular, PDAC is largely refractory to immunotherapy; checkpoint inhibitors are ineffective mainly because of a poor T cell infiltration into the tumor. The TME is largely composed of desmoplastic stroma (forming a physical barrier) and of immunosuppressive cells (i.e., tumor-associated macrophages and neutrophils, myeloid-derived suppressor cells, regulatory T cell) that produce a complex mixture of immunomodulatory effectors exhausting the immune system and preventing its proper functions.^3, 4^ Specifically, NK cells from PDAC patients are functionally altered through the downregulation of activating receptors as well as the reduction of their cytotoxic functions.^5–8^ NK cell immunotherapy has recently become a new important actor in cancer treatment. Compared to T cells, the activation of these cells is based on germline-encoded activating and inhibitory receptors, which therefore does not require any pre-activation to trigger cancer cell death.^9^ Moreover, NK cells can be isolated from a variety of sources and present a safer toxicity profile.^10^ NK immunotherapy encompasses genetically engineered cells such as chimeric-antigen receptor (CAR)-NK cells but also cytokine-based and antibody-based approaches. Importantly, the cytokine IL-15 is well known to stimulate the proliferation and activation of NK cells and superagonists like ALT-803 are currently evaluated in clinical trials for different malignancies.^11, 12^ In PDAC, stimulation of the IL-15/IL-15Rα axis was shown to promote anti-tumor immunity and to enhance the efficacy of PD-1 blockade in vivo.^13^ NK cell engagers (NKCE) are engineered antibody-based structures designed to bind both to NK cells and to tumor cells, in order to stimulate the formation of an immunological synapse between both and consequently trigger specific NK cell activation towards the target.^14^ Various NK cell engagers have been developed and mostly target CD16a (FcγRIIIa), a potent activating receptor that mediates antibody-dependent cellular cytotoxicity (ADCC).^15^ However, limitations to targeting CD16a include polymorphisms leading to decreased binding affinities, downregulation after activation and potential fratricide effects.^16–18^ Other alternative NK cell activating receptor antigens could therefore be used in NK engagers to overcome these hurdles. The co-activation of different NK cell receptors allows enhancing their activation, and synergies between NKp46 and other activating receptors such as 2B4, NKG2D, CD2 and DNAM-1 have been identified.^19^ Similarly, tri-functional constructs targeting both CD16a and NKp46 demonstrated high potency against tumor cells.^20, 21^ Achieving proper NK cell activation by targeting the appropriate NK cell receptors is likely a key element to improve the efficiency of NK cell immunotherapeutic approaches. Growing evidence suggests that increasing NK cell activation, notably through the addition of IL-15 moieties to NK cell engagers, is necessary to properly achievethis.^22–26^ In our present study, we aimed to investigate the efficacy of a combinatorial approach targeting different NK cell activating receptors against PDAC cells. For this, we designed two immunotherapeutic molecules: i) an IL-15Rα/IL-15 based construct associated with anti-NKp46 moieties (called Natural Killer activating Multimeric immunotherapeutic compleXes, NaMiX)^27^ and ii) a Trispecific Killer Engager (TriKE) targeting two of the most expressed activating receptors on NK cells in PDAC, NKG2D and NKp30, and engaging the tumor through the PDAC-associated carcinoembryonic antigen (CEA). In vitro, NaMiX promoted the degranulation of perforin- and granzyme-containing granules in NK cells as well as the production of IFN-γ, thereby enhancing cytotoxicity against pancreatic BxPC-3 cells. NaMiX also increased spheroid cell death, an effect further potentiated by the crosslinking abilities of TriKE that facilitated penetration into the spheroid core and resulted in enhanced death of both spheroids and PDAC patient-derived organoids. Humanized mice bearing BxPC-3 xenografts treated with NaMiX injected 7 times a week (7x/week) exhibited enhanced percentages of human CD45^+^, CD8^+^ T cells and NK cells in the peripheral blood and spleen and a higher NK cell percentage in the tumor. While TriKE tended to delay tumor growth, its combination with NaMiX did not enhance tumor growth inhibition due to the low number of tumor-infiltrating stimulated NK cells in humanized mice. Collectively, these findings underscore the potential of NK cell-based immunotherapeutic approaches for the treatment of PDAC.

## METHODS

Detailed and supplemental methods are described in the supplemental material.

### Cell lines and culture

HEK293F cells (Gibco, #R79007) and HEK293T/17 cells (ATCC, #CRL-11268) were maintained in Dulbecco’s Modified Eagle Medium (DMEM) (Gibco, #11965092) containing 10% heat-inactivated fetal bovine serum (FBS) (Gibco, #10500064), 100 U/ml penicillin + 100 μg/ml streptomycin (Gibco, #15140122) and 2 mM of L-glutamine (Lifetech, #25030024), defined in later sections as “complete DMEM”. Peripheral blood mononuclear cells (PBMCs) were isolated from healthy donors (Red Cross Luxembourg) and maintained in complete Roswell Park Memorial Institute medium (RPMI) (Gibco, #11875093). BxPC-3 cells (pancreatic adenocarcinoma cell line) were purchased from DSMZ (Cat #ACC-760) and cultured in complete RPMI. HT-29 cells (colorectal adenocarcinoma) were purchased from ATCC (Cat #HTB-38) and cultured in complete Iscove’s Modified Dulbecco’s Medium (IMDM) (Gibco, # 12440053). The human NK cell lines NK-92-CD16 (kindly provided by Pr. Béatrice Clemenceau, University of Nantes) and KHYG-1 (DSMZ, #ACC725) were cultured in complete RPMI supplemented with 400 UI/mL rhu-IL-2 (Gentaur, #04-RHUIL-2-3MIU). All cells were cultured at 37 ℃ with 5% CO2.

### Production and molecular characterization of the constructs

The productions and molecular characterization methods are described in the supplemental methods section.

### Binding and crosslinking assays

The specific binding of each construct was confirmed by flow cytometry. BxPC-3 (CEA^+^); NK92-CD16 and KHYG-1 cells (NKG2D^+^ NKp30^+^ NKp46^+^) and HEK293T/17 cells (CEA^-^ NKG2D^-^ NKp30^-^ NKp46^-^) were incubated with 3 μg of the appropriate construct (NaMiX, TriKE or BiKEs) for 30 minutes at 4°C (300,000 cells per condition). After washing, cells were stained with APC-conjugated anti-His Tag antibody. Acquisition was performed on LSR Fortessa flow cytometer (BD Biosciences) and analyzed with Kaluza (Beckman Coulter). The binding assay by ELISA is described in the supplemental methods.

Crosslinking between BiKEs, TriKE and BxPC-3 cells was checked using ELISA by coating a MaxiSorp 96-well flat-bottom ELISA plate (ThermoScientific, # 442404) with His-tagged recombinant CEACAM5 (rCEA) (Invitrogen, #A42584) overnight. Then, 5 μg/mL TriKE or BiKEs was added to the plate, which was revealed with either Fc-tagged recombinant NKG2D (rNKG2D) (Sinobiological, #10575-H01S) or recombinant NKp30 (rNKp30) (Sinobiological, #10480-H02H) followed by goat anti-human Fc-HRP (Invitrogen, #A18817). All incubations with antibodies were done for 1 hour at 4℃, washed using 1% BSA (Carl Roth, #1ET9.1) in PBS (PBS/BSA) and blocked with 5% PBS/BSA. The crosslinking assay by fluorescence and confocal microscopy are described in the supplemental methods section.

### Activation and degranulation assays by ELISA and flow cytometry

Healthy donors PBMCs (Red Cross Luxembourg) (500,000 cells/condition) or PDAC patient PBMCs (BioIVT, # HUMANPBMCCL-0120962) (400,000 cells/condition) were incubated for 48 hours with 3 μg of the molecules or 100 ng human recombinant IL-15 (rhu-IL15) (StemCell Technologies, #78031) or control medium in 100 μl complete RPMI medium in each well of 96 well TC-treated microplates (Sigma Aldrich, #CLS3596-50EA). Control medium (absence of target cells) or 50,000 BxPC-3 and HT-29 cells (presence of target cells) in a volume of 50 μl complete RPMI were added to PBMCs culture, together with anti-CD107a antibody. After 1 hour incubation, GolgiStop (BD Biosciences, #554724) and GolgiPlug (BD Biosciences, #555029) were added for another 4 hours. Cells were centrifuged, washed and stained with LIVE/DEAD Fixable Near-IR Dead Cell Stain Kit (Invitrogen, #L34975), anti-human CD3 (to exclude CD3^+^ T cells), anti-human CD14 and CD19 (to exclude monocytes and B cells), and anti-human CD56 and CD16 to identify NK cells. Cells were fixed, permeabilized following the Cytoperm/Cytofix protocol (BD Biosciences, #554722) and stained with anti-IFN-γ antibody (BD biosciences, #552887). For p-STAT5 and p-rpS6 expression analysis, cells were stained with extracellular staining, fixed in BD Cytofix fixation buffer (BD Biosciences, #554655) and permeabilized using Phosflow Buffer III (BD Biosciences, #558050) according to manufacturer’s instructions before staining with anti-p-STAT5 and anti-p-rpS6 antibodies. For Ki67 expression, the FOXP3/transcription factor staining buffer set was used according to manufacturer’s instructions (Invitrogen, #00-5523-00). Acquisition was performed on the LSR Fortessa flow cytometer (BD Biosciences) and analyzed with Kaluza (Beckman Coulter). Degranulation and cytokine secretion in the supernatant were evaluated using ELISA Flex: Human Granzyme B (HRP) (MabTech, #3486-1H-6), ELISA Flex: Human Perforin (HRP) (MabTech, #3465-1H-6) and ELISA MAX Deluxe Set Human IFN-γ (Biolegend, #430104) following manufacturer’s instructions.

### Cytotoxicity by calcein release assay

The cytotoxicity of PBMCs towards the pancreatic cancer cells was assessed by calcein release assay using a Calcein AM assay kit (Invitrogen, # C34852). Target cells (50,000 cells/condition) were stained following manufacturer’s instructions with 10 μM calcein AM in staining buffer for 30 minutes at 37°C. Stained target cells and effector cells (pre-incubated for 48 hours with NaMiX, rhu-IL15 or control medium) were co-cultured in individual wells of a 96-well microplate at indicated effector:target (E:T) ratios in triplicate for 4 hours at 37°C in a total volume of 200 μl. After incubation, cells were centrifuged and 100 μl of supernatant was harvested in wells of a 96 well Black/Clear Bottom Plates (ThermoScientific, #165305). The fluorescence (F) of the supernatant was measured by a plate reader (Molecular Devices) at Ex/Em = 485/530 nm. The specific lysis was calculated as followed: [F(sample)–F(spontaneous)]/[F(maximum)–F(spontaneous)] x 100%. F(spontaneous) is the fluorescence released from target cells in the absence of effector cells, and F(maximum) represents the fluorescence released after total cell lysis induced by addition of 1% Triton X100 (Sigma-Aldrich, #X100-1L).

### NK cell cytotoxicity assays by flow cytometry

Purified NK cells (300,000 cells/condition) isolated from healthy donors PBMCs (Red Cross Luxembourg) using positive selection with CD56 Microbeads (Miltenyi Biotec,# 130-097-042) or PDAC patient PBMCs (BioIVT, #HUMANPBMCCL-0120962) (400,000 cells/condition) were incubated with NaMiX, rhu-IL15 or control medium for 48 hours similarly to the activation assays described above. After pre-incubation, BxPC-3 cells were stained with CellTrace Violet Cell Proliferation kit (Thermo Fisher Scientific, #C34571) following the manufacturer’s instructions and co-incubated with NK cells for 5 hours at an E:T ratio 1:1 or 2:1 for purified NK cells and 10:1 for PDAC PBMCs. Cells were centrifuged, washed and stained with anti-Annexin V antibody and Propidium iodide (PI) solution (BD Biosciences, #556463) in Annexin V binding buffer (BD Biosciences, #556454). Acquisition was performed on the LSR Fortessa flow cytometer and analyzed with Kaluza. BxPC-3 cells were gated based on CellTrace Violet (BV421) to exclude NK cells and on Annexin V (FITC)/PI (PE) to identify early apoptotic and late apoptotic/other dead cells.

### 3D spheroid cell culture and cytotoxicity assays

To generate 3D spheroids, BxPC-3 cells were stained with CellTrace CFSE dye (Invitrogen, # C34554) and 5,000 CFSE^+^ BxPC-3 cells were seeded in each well of a Nunclon Sphera 96-Well plate (Thermo Scientific, # 174925) and cultured for 48 hours. Healthy donors PBMCs were stained with CellTracker Deep Red (Invitrogen, #C34565) and pre-incubated with NaMiX or control medium for 48 hours. PBMCs were added to BxPC-3 spheroids at an E:T ratio 10:1 in absence or presence of the Engagers and controls including the control scaffold C4BP-β. For cytotoxicity studies, PBMCs or NK cells (isolated from PBMCs) were not stained but were incubated with target cells and molecules in presence of DRAQ7 Deep Red dye (Invitrogen, #D15106). The plates were placed inside the Incucyte S3 (Sartorius) and pictures were acquired every 6 hours over 4 days for kinetics of spheroid’s growth and cytotoxicity. Images were analyzed using Incucyte2024A and ImageJ softwares.

### PDAC organoids culture and functional assay

PDAC organoids were established from a tumor specimen of a patient followed at Strasbourg University Hospital under ethics approval N° CE-2022-49. The participant was informed and did not object to participation in the research or to the use of their data as required by the ethics committe. The study was registered in the public project repository of the Health Data Hub (N° F20220413120650). The protocol for organoid culture was based on Tiriac H et al,^28^ and Broutier et al,^29^ with some modifications. Compositions of the basal medium and PDAC organoid complete medium are described in the supplemental methods section. Briefly, the tumor tissue was collected in 40 mL of IGL-1® organ preservation solution, stored at 4°C and shipped the next day to the Luxembourg Institute of Health. The sample was transferred to a tissue culture dish, the storage buffer was removed, and the sample was mechanically dissociated into fragments of 1 mm^3^ or smaller in the presence of 2 mL of basal medium, on ice. The tissue fragments were then transferred to a Falcon tube and washed twice with 10 mL of basal medium (centrifugation at 200 x g, 5 minutes at 4°C). After the second wash, 10 mL of digestion medium (basal medium containing Gentle collagenase/hyaluronidase (Stemcell tech #7919) 1X and DNase I (Sigma #D5025) at 20 µg/mL) were added to the tube containing the tissue fragments and the enzymatic digestion was conducted at 37°C with a continuous gentle mixing. The digestion was stopped after 2 hours and 40 minutes, the tube was centrifuged at 200 x g for 5 minutes at 4°C, and the supernatant was discarded. The cells were then washed twice with 10 mL of basal medium. After the second wash, all the basal medium was removed carefully, and the tube was placed on ice for 5 minutes. The cell pellet was mixed with 80 µL of Matrigel (Corning #356231) on ice and spotted as a dome in the center of a pre-warmed 12-well plate. After 12 minutes at 37°C, the plate was retrieved from the incubator and 1 mL of pre-warmed PDAC organoid complete medium was dispensed in the well containing the dome. Two days later, several organoids appeared in the Matrigel dome. The medium was changed every 2 to 3 days and, at confluence, the organoids were passaged. To do so, the domes were lifted with a sterile cell lifter and the Matrigel dome was mixed with ice-cold cell recovery solution (CRS) (Corning, #354253) (800 µL to 1 mL/dome) and transferred to a Falcon tube. The depolymerization of the Matrigel was conducted at 4°C in a tube rotator during approximately 40 minutes, followed by a centrifugation at 200 x g for 5 minutes at 4°C. The supernatant was then removed, and the organoid pellet was washed with 10 mL of basal medium. After the washing step, 1 mL of ice-cold basal medium was added to the pellet, and the organoid suspension was mixed several times while hitting the bottom of the tube to help mechanically break up the organoids. The organoid pellet was resuspendend in an adequate volume of Matrigel and spotted as 80-µL domes. For the functional assays, cells within intact organoids (without Matrigel), hereafter referred to as organoid cells, Basal medium was added to bring the volume to 5 mL and the tube was centrifuged. For the functional assays, organoid (without Matrigel) were stained with CellTrace CFSE dye (Invitrogen, # C34554) and 20,000 CFSE^+^ cells were seeded in each well of a Nunclon Sphera 96-Well plate (Thermo Scientific, # 174925).

PBMCs or NK cells (purified from PBMCs) pre-incubated or not with NaMiX for 48 hours were then added to target cells and Engagers in presence of DRAQ7 Deep Red dye (Invitrogen, #D15106) at indicated E:T ratio. The plates were placed inside the Incucyte S3 (Sartorius) and pictures were acquired every 6 hours over 4 days. Images were analyzed using the Incucyte2025B software.

### In vivo treatment of humanized mice bearing BxCP-3 Mice xenografts

Animal experiments were performed in accordance with the local ethics committees and national regulations. The protocol was evaluated by the Animal Welfare Structure of LIH and approved by the Luxembourg Ministry of Agriculture and the Luxembourg Ministry of Health (protocol LUPA 2024/08). The detailed protocol is described in the supplemental methods section. NOD.Cg-Prkdc^scid^ Il2^rgtm1Wjl^/SzJ (NSG) mice were humanized with intravenous injection in the tail vein of 50,000 human CD34^+^ hematopoietic stem cells (HSC) derived from umbilical cord (Lonza, Belgium) in FBS-free RPMI medium. Around 12 weeks post CD34^+^ cells administration, mice were injected with 2,5 μg human recombinant IL-15 (Peprotech, #200-15-10UG) + 7,5 μg of human recombinant IL15Rα (Peprotech, #200-15RA-100UG) intraperitoneally. Around 20 weeks post CD34^+^ cells administration, BxPC-3 cells were injected subcutaneously in the right flank of mice (2 million cells in 100 μl FBS-free RPMI/mouse). Two weeks post-tumor cells engraftment, mice received intraperitoneal injections of the molecules (NaMiX, TriKE, NaMiX + TriKE) at a dose of 1 mg/kg at indicated frequency and control group received PBS every day. Tumor growth was monitored twice a week by Vernier digital caliper. The tumor volume was calculated as follows: Volume = 0.52 x l x w^2^ (l: length of the longest diameter (mm), w=length of the axis perpendicular to l (mm)). At the end of the experiment, mice were sacrificed and peripheral blood, spleen and tumor were excised to isolate cells, which were then analysed by flow cytometry.

### Statistical analysis

Statistical analyses were performed using GraphPad Prism v10 (GraphPad Software, San Diego, CA, USA). All data are presented as individual data with the bars representing mean +/- standard error of the mean (SEM). Comparisons of multiple experiments groups were performed using a one-way analysis of variance (ANOVA), followed by Tukey’s post hoc analysis. For the evaluation of cytotoxicity between the different treatment among different E:T ratios, a two-way ANOVA was used, followed by Tukey’s post hoc analysis. A p-value inferior to 0.05 was considered statistically significant and was indicated as follows: *p < 0.05, **p < 0.01, ***p < 0.001, ****p < 0.0001.

## RESULTS

### Molecular design and binding abilities of NaMiX and Engagers

We have designed two types of immunoconjugates, Natural Killer activating Multimeric immunotherapeutic compleXes (NaMiX) and Trispecific natural Killer Engager (TriKE) to target PDAC cells, and we have compared at first their cytotoxic effects in vitro. NaMiX is composed of the β subunit of the oligomerization domain of the C4 binding protein (C4BP-β) which acts as a matrix for dimerization, as previously reported.^27, 30^ The extracellular sushi domain of IL-15Rα was engrafted on the N-terminal end and single chain variable fragments (scFvs) directed against NKp46, a receptor almost exclusively expressed on NK cells,^21^ were engrafted on the C-terminal end (**Figure 1A**). TriKE was also generated with the C4BP-β scaffold but was embedded in N-terminal with two scFvs targeting the NK cell activating receptors NKG2D and NKp30 (**Figure 1A**). On the C-terminal end of the construct, TriKE contains a camelid single domain antibody (VHH) directed against the tumor-associated antigen CEA, an antigen highly expressed on the surface of different solid tumors, including pancreatic and colorectal cancer.^31, 32^ The structure of TriKE aims to promote the formation of an immunological synapse between both cell types. To compare the effect of the individual and simultaneous NKG2D/NKp30 engagement, we also designed two Bispecific Killer Engagers (BiKEs) targeting only NKG2D (BiKE NKG2D) or only NKp30 (BiKE NKp30) (**Figure 1B**). The molecular patterns of the constructs were analyzed by electrophoretic migration and Western Blot (**Supplemental Figure 1A**). Under non-reducing (NR) conditions, NaMiX shows mostly a band around 110 kDa representing the dimeric form, which disappears in favor of the monomeric form around 50 kDa under reducing (R) conditions. Similar patterns were observed for TriKE and BiKEs, indicating that all constructs are produced primarily as dimers. After selection and expansion, the selected clones were purified with nickel-based affinity chromatography (**Supplemental Figure 1B**). A second chromatography was performed to purify TriKE using TwinStrepXT affinity matrix to ensure the specific isolation of the heterodimers, containing both NKG2D and NKp30 binding moieties. The binding specificity of all constructs towards their target was then evaluated by flow cytometry (**Supplemental Figure 1C-F**). When incubated with NaMiX, an increased percentage of His Tag-positive cells was observed for a NKp46^+^ cell line (NK92-CD16 cells) but not for a NKp46^-^ cell line (HEK cells) (**Supplemental Figure 1C**). TriKE and BiKEs showed binding to the CEA^+^ BxPC-3 cell line and to the KHYG-1 cell line expressing NKG2D and NKp30 but not to HEK-293T cells (**Supplemental Figure D-F**). The specificity of binding of each construct was further confirmed by competition ELISA with recombinant NKp46 for NaMiX, and with recombinant NKG2D, NKp30 or CEA for the TriKE and BiKEs (**Supplemental Figure 1G**).

**Figure 1:**
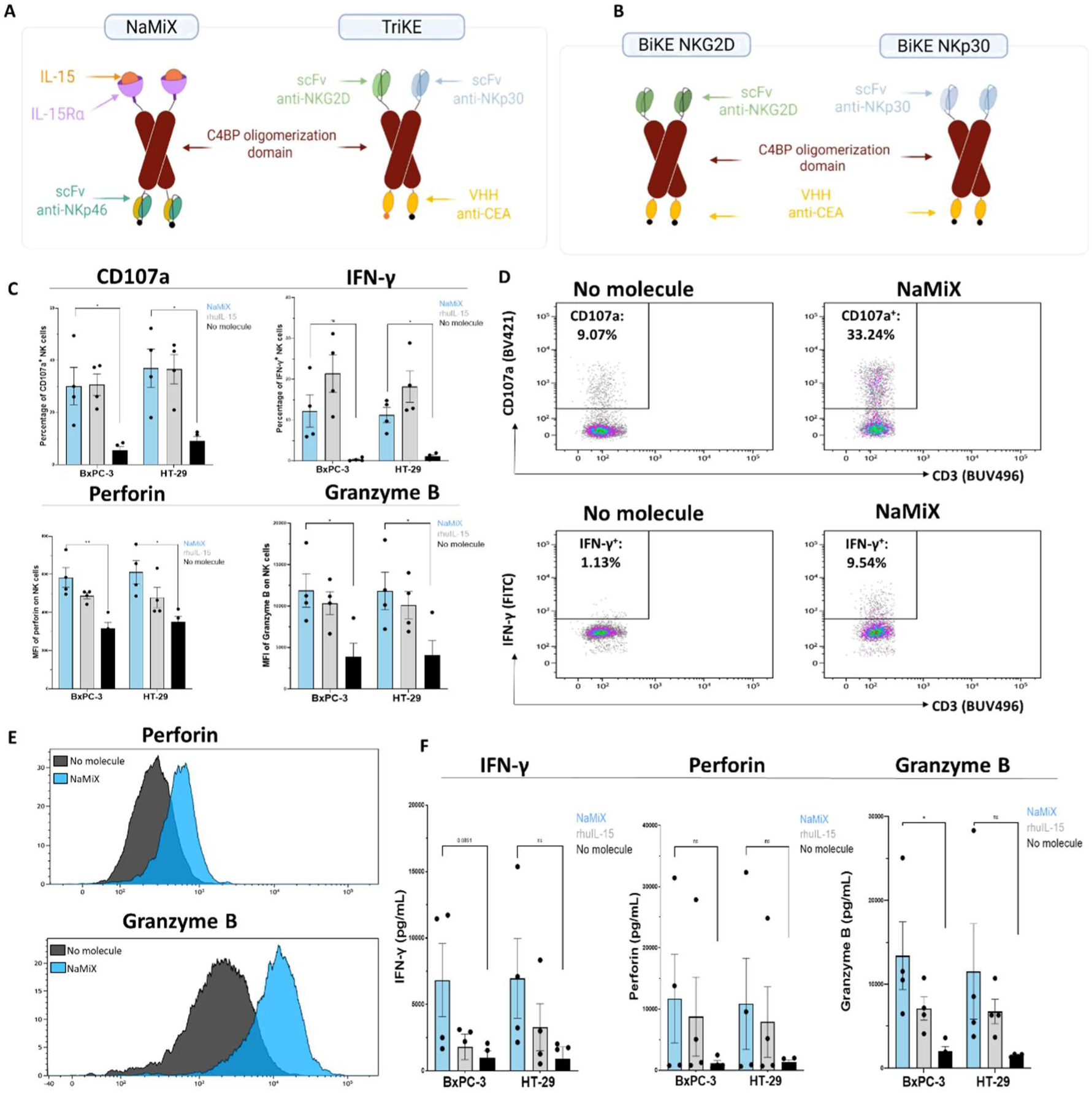
NaMiX stimulates the activation and degranulation of PBMCs against the pancreatic cancer cell line BxPC-3 and the colorectal cancer cell line HT-29. (A-B) Both NaMiX and TriKE expressed dimers upstream and downstream of the dimerization domain of the C4 binding protein β (C4BP-β). NaMiX is composed of IL-15Rα/IL-15 complexes associated to anti-NKp46 single chain variable fragments (scFvs) and TriKE and BiKEs are composed of scFvs targeted against NKG2D and/or NKp30 and VHH against CEA. PBMCs were pre-incubated with NaMiX, rhu-IL15 or control medium for 48 hours, and BxPC-3 cells were added for 5 additional hours at an effector:target ratio of 10:1. (C) NK cells were then analyzed by flow cytometry for their expression of CD107a, IFN-γ, perforin and granzyme B. (D-E) Representative dot plots for CD107a and IFN-γ expression and histograms for perforin and granzyme B expression. (F) The cell culture supernatant was analyzed for perforin, granzyme B and IFN-γ by ELISA. Data are presented as the mean values ± SEM. Results correspond to two pooled independent experiments (2 donors per experiment). Statistical analysis was performed using a one-way ANOVA and post-hoc Tukey test (*p<0.05; **p<0.01).

### NaMiX stimulates the activation and degranulation of PBMCs against pancreatic and colorectal cancer cell lines

We then confirmed that NaMiX could stimulate NK cell activation and degranulation on PBMCs of healthy donors (HD) similarly to the original NaMiX targeting NKG2A and KIR2DL (**Supplemental Figure 2**).^27^ After 48 hours of incubation, NaMiX significantly increased extracellular CD107a and intracellular IFN-γ (p<0.01) expression in NK cells (**Supplemental Figure 2B**). Higher levels of IFN-γ (p<0.01) and granzyme B (p<0.001) were released in the cell culture supernatant (**Supplemental Figure 2C**). Perforin expression was enhanced both intracellularly and in the supernatant, although non-significantly. We further assessed whether NK cell activation-mediated by NaMiX was associated with the activation of IL-15 dependent pathways in NK cells, the STAT3/5-JAK and Akt-mTOR pathways.^33^ Upon short-term exposition (5 minutes), NaMiX induced the phosphorylation of STAT5, but not of rpS6, as opposed to rhu-IL15 (**Supplemental Figure 2D**). Concordantly to the role of mTOR in NK cell proliferation, NaMiX only stimulated a low increase in proliferation (24,7 % of Ki67^+^ cells) as compared to the control medium (3,11 % of Ki67^+^ cells) and rhu-IL15 (69,95 % of Ki67^+^ cells), respectively (**Supplemental Figure 2E**). Finally, we observed a significant upregulation of NKp30 expression by two fold (p<0.05) but not of NKG2D (p=0.77) upon a 48h exposition to NaMiX (**Supplemental Figure 2F**). These results show that NaMiX stimulates the degranulation and activation but only low proliferation of NK cells. Next, we evaluated whether NaMiX was able to trigger NK cell activation towards the tumor-associated antigen CEA^+^ cancer cell lines BxPC-3 (pancreatic ductal adenocarcinoma cells) and HT-29 (colorectal adenocarcinoma cells). In presence of BxPC-3 and HT-29 cells, NK cells incubated with NaMiX showed increased degranulation compared to the control medium as evidenced by increased CD107a, perforin and granzyme B expression (p<0.05) (**Figure 1C-E**). The percentage of IFN-γ^+^ NK cells was also upregulated, although non-significantly against BxPC-3 cells (p=0.09). Furthermore, this increased degranulation was also observed by ELISA with a higher concentration of IFN-γ, perforin and granzyme B upon NaMiX incubation, although the high variability between donors did not allow reaching statistical significance (**Figure 1F**).

### **N**aMiX stimulates cytotoxicity of PBMCs and purified NK cells against K562 and pancreatic cancer cell lines

We then assessed whether pre-incubation of PBMCs with NaMiX was able to stimulate cytotoxicity against cancer cells by a calcein release assay (**Figure 2A-B**). We observed an increased cytotoxicity of PBMCs incubated with NaMiX towards BxPC-3 cells, and this effect was proportional to the effector:target (E:T) ratio (**Figure 2A**). We then compared these results obtained with the cytotoxicity triggered to K562 E:T (ratio of 10:1) as gold standard for assessing NK cell cytotoxicity, since these cells lack HLA-I expression, and HT-29, another CEA^+^ cell line known to be highly resistant to NK cell action due to their high HLA-E expression levels (**Figure 2B)**.^34^ We observed that NaMiX induced similar cytotoxicity towards the K562 cell line (70.33 %, p<0.01) but was less efficient towards HT-29 cells (40.67 %, p=0.08), suggesting a main cytotoxic role of NK cells in this effect. To confirm this hypothesis, we isolated NK cells from PBMCs and evaluated their cytotoxicity against BxPC-3 cells. NaMiX stimulated the NK cells-mediated cytotoxicity of target cells, as evidenced by an increase of early apoptotic cells (Annexin V^+^ PI^-^) (although non-significant) and a significant increase in the percentage of cells undergoing late regulated cell death mechanisms (Annexin V^+^ PI^+^) (p<0.01) (**Figure 2C-D**). Importantly, NaMiX was also able to trigger activation and cytotoxicity of PBMCs isolated from a PDAC patient toward BxPC-3 cells (**Figure 2E)**. Collectively, these results show that NaMiX stimulates the cytotoxicity of PBMCs, of both healthy donors and a PDAC patient, and purified NK cells against various cancer cell lines, including pancreatic cancer BxPC-3 cells.

**Figure 2:**
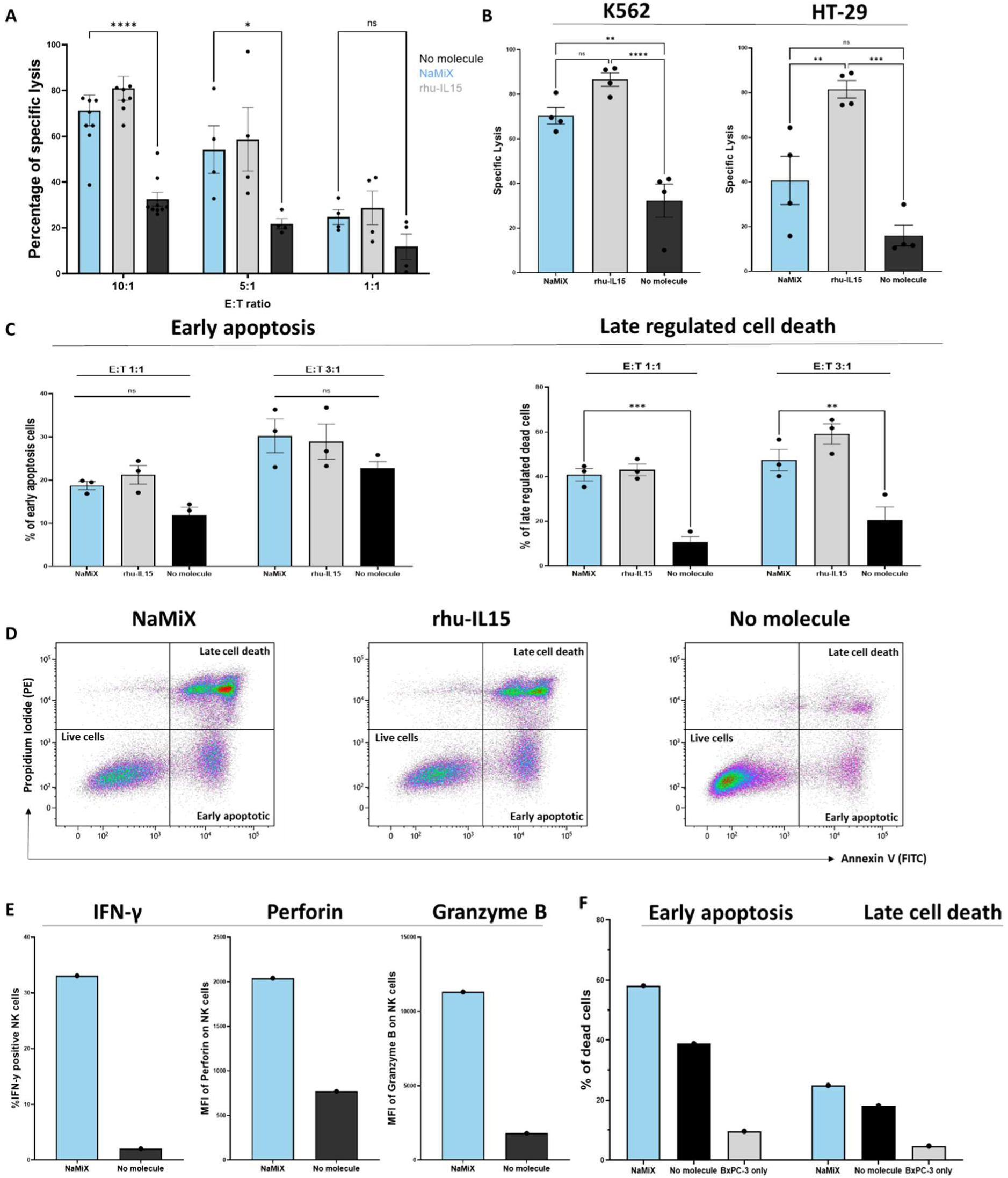
NaMiX stimulates the cytotoxicity of PBMCs and purified NK cells against the pancreatic cancer cell line BxPC-3, the colorectal cancer cell line HT-29 and the myeloblast cell line K562 at different effector:target ratios. The cytotoxicity of PBMCs or purified NK cells against target cells was assessed by (A-B) calcein release assay and (C-D) flow cytometry. For the calcein release assay, BxPC-3 (A), K562 or HT-29 (B) were stained with calcein AM and incubated for 5 hours with PBMCs (pre-activated for 48 hours with NaMiX, rhu-IL15 or control medium) at effector:target (E:T) ratio of 10:1 unless stated otherwise. The fluorescence of the supernatant was quantified to calculate the percentage of specific lysis. For the flow cytometry assay, BxPC-3 cells were stained with CellTrace Violet and incubated for 5 hours with NK cells purified from PBMCs (pre-activated for 48 hours with NaMiX, rhu-IL15 or control medium) at indicated E:T ratios. An Annexin V/Propidium iodide (PI) staining was performed and analyzed as follows: viable BxPC-3 cells appear as CellTrace Violet^+^/Annexin V^-^/PI^-^, early apoptotic BxPC-3 cells appear as CellTrace Violet^+^/Annexin V^+^/PI^-^ and BxPC-3 cells undergoing late cell death and other regulated cell death mechanisms appear as CellTrace Violet^+^/Annexin V^+^/PI^+^. Results correspond to two pooled independent experiments (1-4 donors per experiment). Statistical analysis was performed using a two-way ANOVA and post-hoc Tukey test (*p<0.05; **p<0.01; ***p<0.001; ****p<0.0001). (E-F) PBMCs isolated from one PDAC patient were pre-activated or not with NaMiX and co-incubated with BxPC-3 cells; (E) Intracellular expression of IFN-γ, perforin and granzyme B was evaluated on NK cells and (F) Mortality of BxPC-3 cells was evaluated by AnnexinV/PI staining. Data are presented as the mean values ± SEM.

### NKp30 Engagers increase NaMiX-mediated NK cell cytotoxicity against BxPC-3 spheroids by crosslinking

To complement our first cytotoxic approach, we next aimed to evaluate the effect of a trispecific killer Engager (TriKE) targeting NKG2D and NKp30 as well as CEA, a tumor-associated antigen (TAA) expressed by PDAC cells, to engage NK cells near BxPC-3 cells. We first demonstrated the ability of our TriKE to bind simultaneously to NK cell and cancer cell antigens by ELISA (**Figure 3A-B**). Similar results were obtained with the BiKEs targeting only NKG2D (BiKE NKG2D) or NKp30 (BiKE NKp30) (**Supplemental Figure 3A**), demonstrating that all Engagers are able to simultaneously and specifically bind to CEA and targeted NK activating receptors antigens. Fluorescence microscopy and confocal microscopy analyses confirmed that TriKE stimulated the formation of immunological synapses between the NK cell line KHYG-1 and pancreatic cancer BxPC-3 cells (**Figure 3C-D**). We next evaluated whether NaMiX and Engagers could improve the cytotoxicity of PBMCs and NK cells against PDAC targets in 3D cellular models. Alone, the simultaneous co-engagement of NKG2D and NKp30 by TriKE seemed to increase the cytotoxic activity of PBMCs against BxPC-3 spheroids, as evidenced by a trending increase of IFN-γ production (**Supplemental Figure 3B)** and a trending decreased spheroid size after 4 days of co-culture, although these effects were highly donor-dependent (**Supplemental Figure 3C-D).** Interestingly, the combination of NaMiX with anti-NKp30 constructs (both BiKE NKp30 and TriKE) enhanced the PBMCs’ cytotoxicity towards BxPC-3 spheroids starting from 48 hours of co-incubation (p<0.01), an effect that is not observed with the combination of NaMiX and BiKE NKG2D or control scaffold C4BP-β (**Figure 3E)**. This effect was not enhanced by targeting both NKp30 and NKG2D (TriKE) as compared to NKp30 alone (BiKE NKp30). This potentiated NKp30 effect led to almost complete destruction of BxPC-3 spheroids after 72 hours of co-incubation (**Figure 3F-G)**. Further, spheroids incubated with purified NK cells stimulated by NaMiX and NKp30-targeting Engagers exhibited an intense staining of the cytotoxicity marker DRAQ7, which co-localized with the spheroid core (**Figure 3H).** Similar results were obtained with PBMCs isolated from a PDAC patient (**Figure 3I**). Altogether, these results demonstrate that the combination of NaMiX and NKp30 Engager enhances PBMCs and NK cells cytotoxicity towards BxPC-3 spheroids.

**Figure 3:**
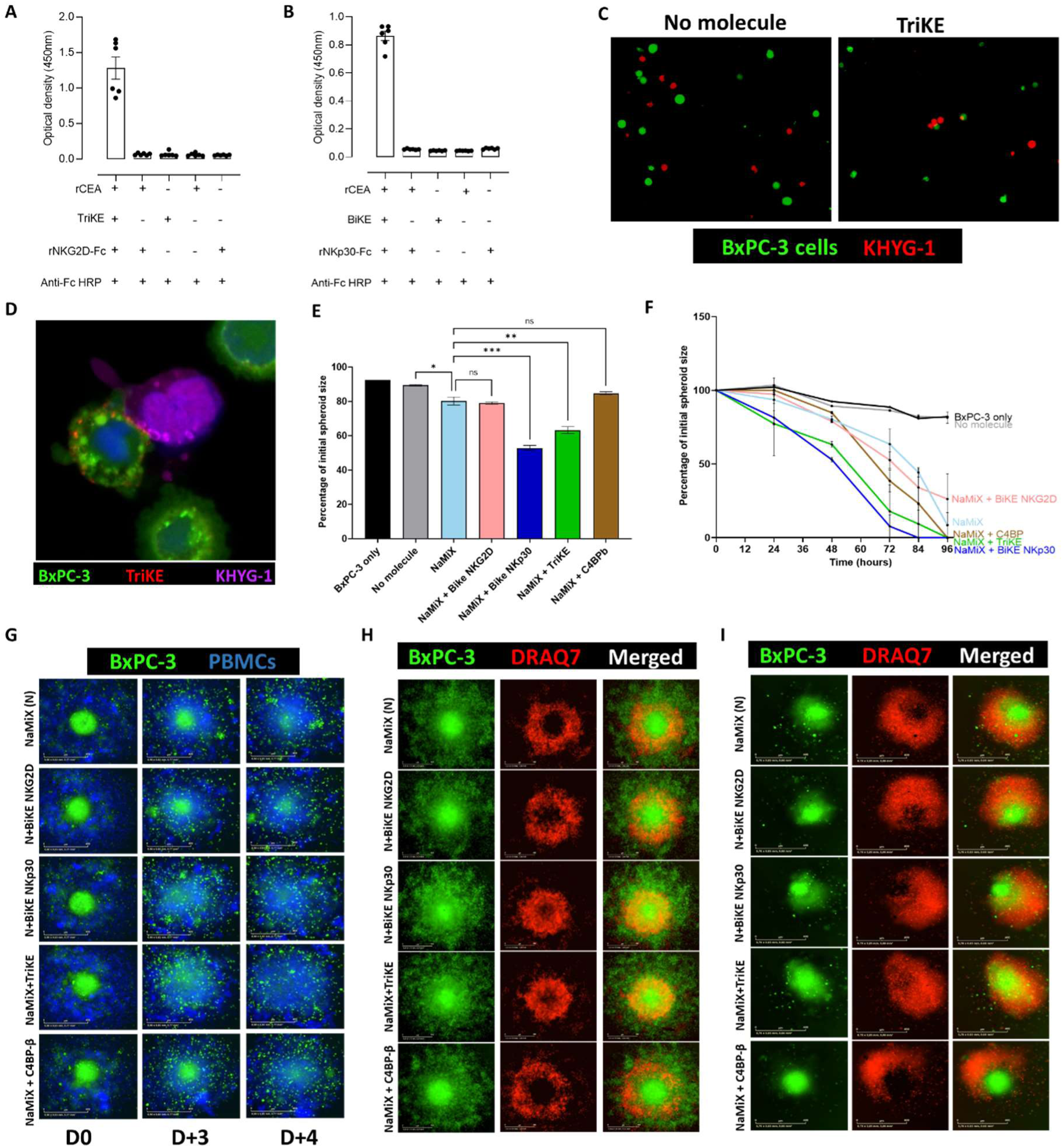
NKp30 engagers increase NaMiX-mediated NK cell cytotoxicity against BxPC-3 spheroids by increasing the cross-linking between both cell types. (A-B) ELISA assay using recombinant human CEA (rCEA) as coating, adding TriKE, then revealing by (A) recombinant NKG2D coupled to Fc (rNKG2D-Fc) or (B) recombinant NKp30 coupled to Fc (rNKp30-Fc) and an anti-Fc linked with HRP. (C-D) CFSE-stained BxPC-3 (green) and CellTracker Deep Red-stained KHYG-1 (red/pink) cells were co-cultured with (C) control medium or with TriKE and analyzed by fluorescence microscopy or (D) with AlexaFluor647-labeled TriKE (red) and analyzed by confocal microscopy. (E-G) CFSE-stained BxPC-3 were seeded in ultra-low attachment plates for 48 hours to form spheroids, then co-incubated with CellTracker Deep Red stained PBMCs pre-incubated with NaMiX or control medium for 48 hours, together with BiKEs, TriKE, control scaffold or control medium and placed inside IncucyteS3 for 96 hours. Spheroid size was assessed using ImageJ software and the percentage of initial spheroid size is represented (E) after 48 hours of co-incubation or (F) over 96 hours of co-incubation. (G) Representative image at day 0, day 3 and day 4 of co-incubation. (H-I) Representative images of CFSE-stained BxPC-3 co-incubated with purified NK cells from a healthy donor (H) and a PDAC patient (I) and the cytotoxicity marker DRAQ7 at day 3 of co-incubation with indicated molecules. Data are presented as the mean values ± SEM. Results correspond to 2 pooled representative donors. Statistical analysis was performed using a one-way ANOVA and post-hoc Tukey test (*p<0.05; **p<0.01; ***p<0,001). Scale bar = 400 µm.

### The combination of NaMiX and NKp30 Engagers trigger higher NK cell cytotoxicity against patient-derived PDAC organoids

To complement our models, we evaluated the combination of NaMiX and Engagers on NK cell cytotoxicity towards patient-derived organoids (PDOs) (**Figure 4**). While NaMiX enhanced the cytotoxic activity of purified NK cells against PDOs, the combination of NaMiX and Engagers (particularly BiKE NKp30 and TriKE) resulted in a further increase in cytotoxicity. This is evidenced by an enhanced DRAQ7 staining and a marked reduction in organoid initial size that reached 37.47% of initial size after 4 days of co-culture with NaMiX + TriKE (**Figure 4A-C**). This effect was proportional to the E:T ratio, and was also observed with PBMCs (**Supplemental Figure 4)**. Altogether, NK cells activated by NaMiX and engaged through the NKp30 receptors exhibit higher cytotoxic potential towards pancreatic cancer cells in 3D models.

**Figure 4:**
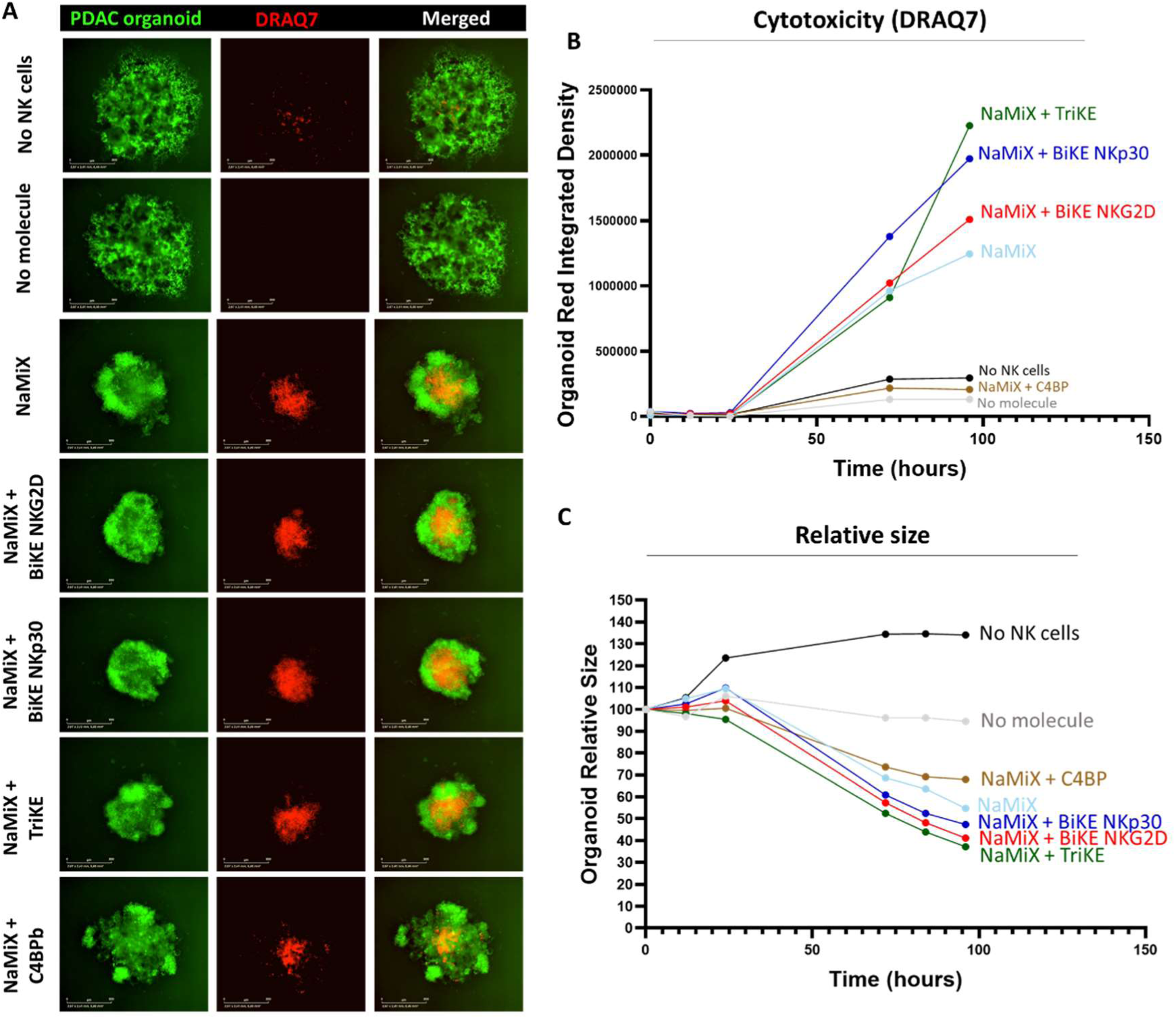
The combination of NaMiX and NKp30 Engagers triggers higher NK cell cytotoxicity against patient-derived PDAC organoids. CFSE-stained patient-derived organoids were seeded in ultra-low attachment plates together with purified NK cells and with the cytotoxicity marker DRAQ7 with indicated Engagers in duplicates and placed inside IncucyteS3 for 96 hours. (A) Representative images at day 4 of co-incubation. Incucyte 2025B software was used to quantify (B) the red integrated density and (C) and organoid relative size. The experiment was performed 3 times and one representative donor is shown in A, B, C. Scale bar = 800 µm.

### NaMiX enhances the development of cytotoxic lymphocytes in humanized mice bearing BxPC-3 xenografts

Furthermore, we developed a humanized mice model bearing subcutaneous BxPC-3 xenografts to evaluate the in vivo effect of our immunoconjugates on human NK cells development, activation and antitumoral cytotoxicity **(Figure 5A**). Mice humanization was performed through the intravenous administration of umbilical-cord derived CD34^+^ hematopoietic stem cells (HSCs), and resulted in the development of the main human immune cell populations after 18 weeks **(Supplemental Figure 5**). Since NK cell activation in vitro is solely supported by NaMiX in our dual strategy, we evaluated the effects of NaMiX alone on NK cell development according to two different doses per week. NaMiX was therefore administered intraperitoneally (IP) 3 times (3x/week) or 7 times (7x/week) over one week and the percentage of immune cell populations was quantified in the peripheral blood, spleen and tumor xenograft. When administered every day (7x/week), NaMiX stimulated significantly the expansion of human CD45 (hCD45) in the tumor (p<0.05) and blood (p<0.01), as well as increased the percentage of human NK cells relative to all CD45 in the tumor (8.37% vs 2.53%, p<0.05), and strongly in the blood (11.29% vs 2.09%, p<0.01) and spleen (15.16% vs 3.65%, p<0.0001) as compared to the PBS control. It was a higher effect than for the condition NaMiX injected 3x/week (**Figure 5B-C**). In the tumor, NaMiX 7x/week increased the percentage of tumor-infiltrating NK cells relative to all xenograft cells from 0.52% with PBS mice to 2.44% (p<0.001) (**Figure 5F**). Interestingly, NaMiX treatment also significantly triggered the development of other cytotoxic lymphocytes such as CD8^+^ T cells in the blood and spleen (**Figure 5D**) and Natural Killer T (NKT) cells in the tumor and spleen (**Figure 5E**). However, NaMiX did not have any impact on other immune cell populations including CD19^+^ B cells, CD14^+^ monocytes and dendritic cells (DC), as shown on the t-stochastic neighbour embedding (t-sne) analysis of splenic immune cell populations (**Figure 5G**).

**Figure 5:**
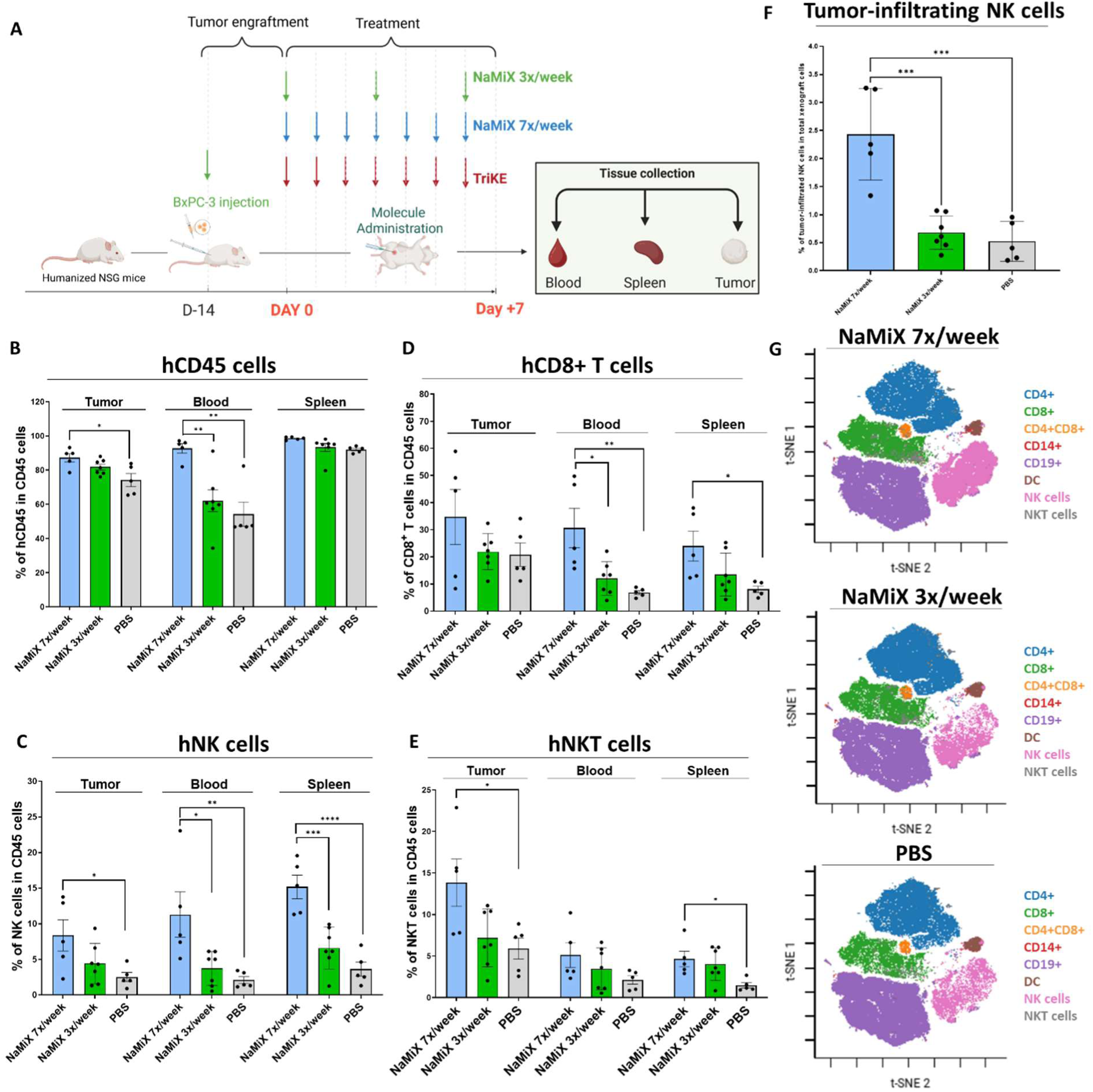
NaMiX stimulates the development of cytotoxic lymphocytes in humanized mice bearing BxPC-3 xenografts. 20 weeks-old humanized NSG mice were subcutaneously injected 2 million of BxPC-3 cells and intraperitoneally administered with 1 mg/kg of indicated constructs. Mice received treatment for 1 week with NaMiX given every 3 days (NaMiX 3x/week) or every day (NaMiX 7x/week), and TriKE and PBS given every day. (A) Schematic representation of the in vivo experiment. (B-F) After one week of treatment, tumor, blood and spleen were excised and the percentage of (B) hCD45, (C) NK cells, (D) CD8^+^ T cells and (E) NKT cells was analyzed by flow cytometry and expressed as proportion from all CD45 cells. (F) Percentage of tumor-infiltrating NK cells relative to all xenograft cells. (G) tSNE plots of the clustering of splenic immune cell populations in the different groups were generated. Data are presented as the mean values ± SEM (5-7 mice per group) Statistical analysis was performed using a one-way ANOVA and post-hoc Tukey test (*p<0.05; **p<0.01; ***p<0.001; ****p<0.0001).

### NaMiX induces the remodeling of the cytotoxic NK cells repertoire in vivo

Finally, we evaluated whether NaMiX could sustain its anti-tumoral effect in vivo against BxPC-3 xenografts, and whether this effect was potentiated with TriKE, similarly to what was observed in spheroids and PDOs. A slight tumor growth delay (p=0.1583) was observed in mice treated with TriKE as compared to PBS, suggesting a beneficial effect of the crosslinking between NK cells and PDAC cells. However, NaMiX did not result in significant tumor growth delay in mice when given 3x/week or 7x/week, and this effect could not be improved in combination with TriKE (**Figure 6A**). To explain this, we first assessed the functional and phenotypic characteristics of splenic NK cells, as this population is abundant and should be more exposed to NaMiX than infiltrated NK cells. Interestingly, splenic NK cells from NaMiX 7x/week-treated mice and re-stimulated ex vivo with BxPC-3 cells exhibited significantly higher intracellular content of granzyme B (p<0.001) and perforin (p<0.05), as well as increased IFN-γ expression (p=0.1087) as compared to PBS mice, and this effect was not improved by combination with TriKE (**Figure 6B-C**). This data supports that NaMiX treatment, independently from TriKE, mediates NK cells functional activation in vivo. However, expression of CD107a was not enhanced upon NaMiX treatment resulting in a non-increase of cytotoxicity towards BxPC-3 cells ex vivo as compared to the PBS control (**Figure 6D-E**). This indicates that despite an accumulation of cytotoxic granules within NK cells upon treatment, NaMiX could not stimulate the degranulation and cytotoxicity of NK cells towards BxPC-3, both in vivo and ex vivo. To understand the mechanisms of potential tumor resistance upon NaMiX treatment in this model, dimensional reduction was conducted after flow cytometry based on the expression of immune cell lineage markers (CD3, CD8, CD14/CD19, CD16, CD56) and 10 clusters were defined, among which 3 clusters were relative to NK cells (Clusters 2, 3 and 7) (**Figure 6F**). The relative abundance of Cluster 2 (CD56^dim^CD16^+^), Cluster 3 (CD56^+^CD16^dim/-^) and Cluster 7 (CD56^-^CD16^+^) was then quantified among treatment groups. Cluster 2, which displays a phenotype consistent with cytotoxic NK cells, was upregulated and became the predominant NK cell subset in NaMiX 7x/week compared to PBS control. An increase in Cluster 3, associated with a CD56^dim^ regulatory phenotype, was also observed whereas Cluster 7 did not show any increase (**Figure 6G**). We further analyzed the expression of the main inhibitory NK cell receptors (CD158e1, NKG2A, CD158ah and TIM-3) on each of these clusters among the treatment groups. Notably, repeated NaMiX administrations led to upregulation of these receptors on Cluster 2, and to a lesser extent Cluster 3, suggesting a negative retro-control loop that might inhibit NK cell functions following the production of granzyme and perforin-containing granules (**Figure 6H**). However, expression of the immune checkpoints PD-1 and TIGIT remained unchanged on both splenic NK cells and CD8^+^ T cells, indicating that this inhibitory state likely does not represent an exhausted phenotype (**Figure 7A)**. Importantly, while the low quantity of intra-tumoral NK cells precluded functional analyses, phenotypic characterization of the intratumoral cytotoxic CD56^dim^CD16^+^ NK cell subset showed that NaMiX treatment induced a less inhibited phenotype compared with splenic NK cells. This was evidenced by a moderate upregulation of NKG2A, TIM-3 and inhibitory KIRs alongside increased expression of the activating receptor NKG2C and a decreased expression of the inhibitory receptor KLRG1 (**Figure 7B**). Taken together, these results highlight that while repeated NaMiX administration might lead to peripheral NK cell inhibition following activation, a more activated anti-tumor phenotype of intratumoral NK cells is observed, although NK cells infiltration was too low to enable tumor regression.

**Figure 6:**
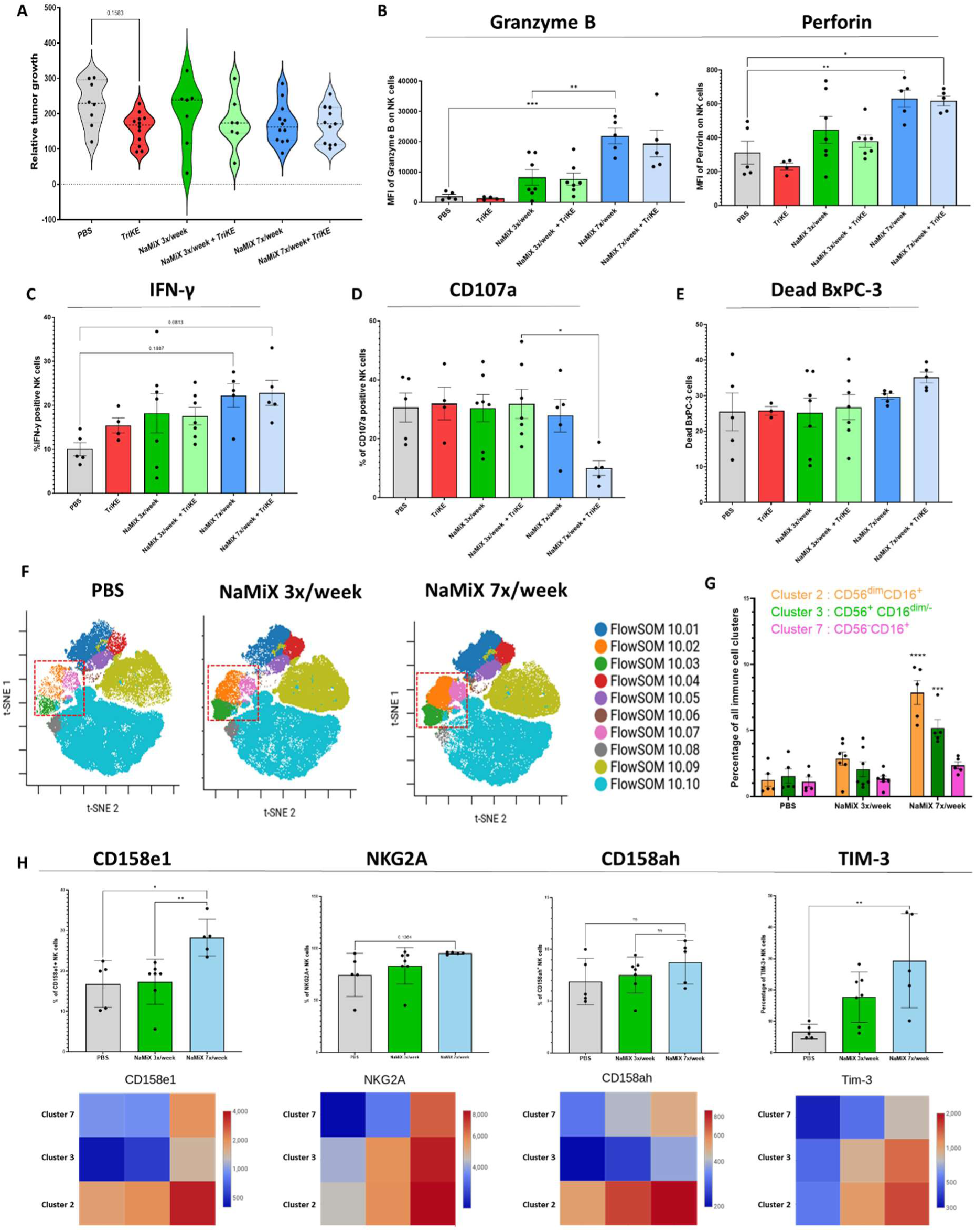
In vivo evaluation of NaMiX and TriKE-mediated activation of NK cells. 20 weeks-old humanized NSG mice were subcutaneously injected 2 million of BxPC-3 cells and intraperitoneally administered with 1 mg/kg of indicated constructs. Mice received treatment for 1 week with NaMiX given every 3 days (NaMiX 3x/week) or every day (NaMiX 7x/week); TriKE and PBS were given every day. (A) Tumor size was measured after one week of treatment and the relative size was calculated as the percentage from initial tumor size. (B-D) Splenic cells from treated mice were incubated ex vivo with BxPC-3 cells for 5 hours then analyzed for their expression of (B) granzyme B, perforin, (C) IFN-γ and (D) CD107a. (E) BxPC-3 mortality was assessed by Live/Dead staining. (F) T-sne plots from dimensional reduction based on the expression of immune cell lineage markers with 10 defined clusters. (G) Relative abundance of each cluster among treatment groups. (H) Percentage (histogram) and relative expression (heat map) of CD158e1, NKG2A, CD158ah and TIM-3 of NK cells among treatment groups. Data are presented as the mean values ± SEM (7-12 mice per group for tumor growth; 5-7 mice per group for other analyses). Statistical analysis was performed using a one-way ANOVA and post-hoc Tukey test (*p<0.05; **p<0.01; ***p<0,001).

**Figure 7:**
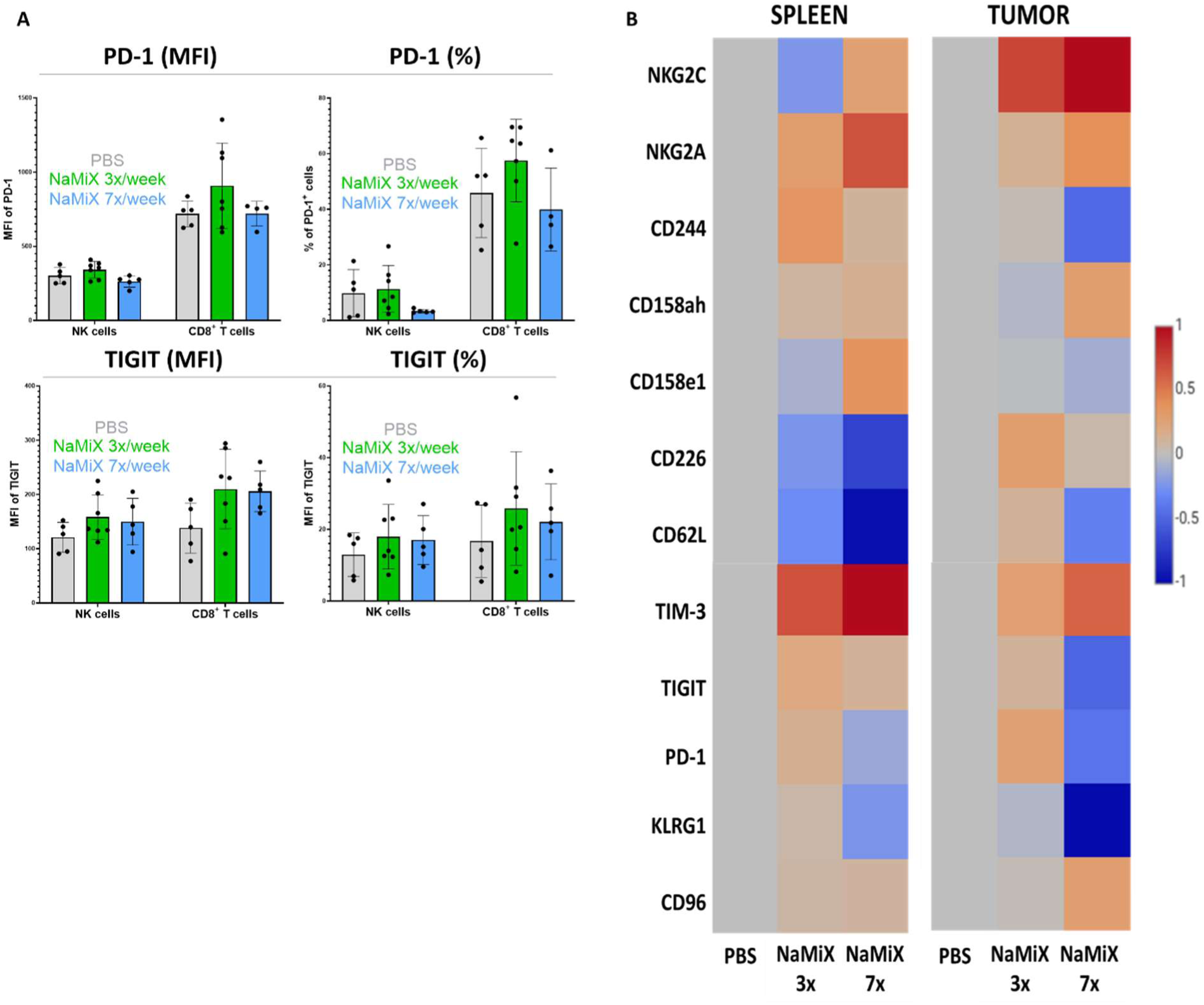
Phenotypic characterization of NK cells in treated humanized mice. 20 weeks-old humanized NSG mice were subcutaneously injected 2 million of BxPC-3 cells and intraperitoneally administered with 1 mg/kg of indicated constructs. Mice received treatment for 1 week with NaMiX given every 3 days (NaMiX 3x/week) or every day (NaMiX 7x/week), PBS was given every day. (A) MFI and percentage of PD-1 and TIGIT on splenic NK cells and CD8^+^ T cells among treatment groups. Data are presented as the mean values ± SEM (5-7 mice per group). Statistical analysis was performed using a one-way ANOVA and post-hoc Tukey test. (B) Heat maps of relative expression of NK cell receptors of treated mice compared to PBS group in spleen and tumor.

## DISCUSSION

Pancreatic ductal adenocarcinoma represents one of the most challenging cancer types, and new therapeutic approaches are urgently needed to improve the prognosis of the patients. As such, NK cells immunotherapy is becoming an increasing new actor in cancer immunotherapy, and their stimulation and engagement were investigated here against PDAC using 2D and 3D cellular models as well as humanized mice bearing functional human NK cells. To boost NK cell function against their targets, one strategy consists in the administration of recombinant human IL-15 (rhuIL-15), a cytokine known to stimulate the activation and proliferation of NK cells.^33^ Although rhuIL-15 is able to stimulate NK cell functions in vivo, the very short half-life of this cytokine requires repetitive administration to maintain an efficient therapeutic concentration, leading to toxic side effects.^35^ The development of IL-15 superagonists has therefore become an important topic of research. In a first step, we evaluated NaMiX, an immunoconjugate possessing IL-15/IL-15Rα complexes associated with anti-NKp46 moieties to target NK cells. NKp46 is often considered the most specific NK cell receptor, and its expression is consistent among NK cell subtypes and activation states.^16, 17^ Moreover, the stimulation of NKp46-related signaling pathways leads to the phosphorylation of ZAP70/Syk and subsequent activation of the PI3K pathway, associated with NK cell activation and cytotoxicity.^36^ NKp46 expression remains high in PDAC patients, making it an interesting target for our approach,^5^ since we have previously shown that the level of receptor expression on NK cells is crucial for proper efficiency of NaMiX.^27^ Cancer cells and the associated tumor microenvironment (TME) are known to inhibit NK cell functions, notably by downregulating activating receptors and impairing degranulation.^37–39^ Still, in presence of target cells expressing CEA, we showed that NaMiX specifically targets and activates NK cells in vitro through binding to NKp46 and stimulation of IL-15 signaling pathway resulting in the release of more cytotoxic perforin- and granzyme B-containing granules, in particular when using PBMCs from PDAC patients. Further, NaMiX induced sufficient NK cell degranulation and activation to trigger significant cytotoxicity towards pancreatic cancer cells BxPC-3 in 2D and 3D models in vitro. Accessing pancreatic cancer cells is challenging due to the hostile tumor microenvironment surrounding the tumor.^3^ Therefore, we investigated in a second step whether the engagement between NK cells and pancreatic cancer cells through NKCE targeting activating receptors could further engage and enhance the effects of NaMiX. NKCE represent a growing field, and a number of constructs have been reported mainly for the treatment of liquid tumors and few for some solid tumors.^14^ To our knowledge, no NKCE has been reported for the treatment of PDAC. CD16a is a popular target for NK cell engagers, due to its ability to mediate ADCC, and the synergy between CD16a and other activating receptors such as NKp46 and NKG2D has already been described.^19^ Here, we developed a tri-specific killer Engager (TriKE) with scFvs against NKG2D and NKp30 since NKG2D and NKp30 are still amongst the most expressed receptors in PDAC patients,^5^ and persistent NKp30 expression has been described, even after NK cell activation.^17^ NK cell engagers targeting NKp30 through anti-NKp30 fragments or through its ligand B7-H6 exhibited efficacy against multiple myeloma and EGFR-expressing cancer.^40, 41^ Moreover, a few NKG2D targeting engagers have also been reported, and multiple clinical trials are ongoing to evaluate the efficacy of CD16a x NKG2D constructs associated with anti-HER2, EGFR, CD33 and BCMA in locally advanced or metastatic solid tumors (NCT04143711; NCT05597839; NCT04789655; NCT04975399).^42–45^ Aside from its ability to trigger rapid NK cell activation, NKG2D is also expressed in other cell types including NKT cells, a subset of CD8^+^ T cells and a subset of γδ T cells, which can provide additional cytotoxic effects against the target. We showed that the combination of anti-NKp30 Engagers (TriKE and BiKE NKp30) with NaMiX significantly improved NK cells cytotoxicity towards BxPC-3 spheroids and patient-derived organoids. The high cytotoxic action of NaMiX coupled to the crosslinking abilities of the Engagers allowed NK cells to reach and attack the core of the spheroid, resulting in spheroid destruction after 3-4 days of co-culture. In contrast, the combination of NaMiX and BiKE NKG2D did not enable to increase the spheroid destruction. This might be partially explained by the increased NK cells expression of NKp30 (but not NKG2D) induced by NaMiX pre-incubation. Additionally, targeting NKG2D can be challenging, as a desensitization of this receptor after chronic engagement leading to impaired effector functions has been reported.^46^

Humanized mice models have become important translational tools in immuno-oncology research.^47^ Using CD34^+^ HSC transplanted NSG mice with a single administration of IL-15 and IL-15Rα to promote the proliferation of NK cells,^48^ our model demonstrated the development of the main human immune cell populations, including CD19^+^ B cells, CD3^+^ T cells (both CD4^+^ and CD8^+^), NK cells and CD14^+^ monocytes. Our humanized mouse model is designed to closely recapitulate the human immune system, allowing investigation of complex interactions among distinct immune cell subsets after establishment of subcutaneous BxPC-3 xenografts. This is critical, as NK cell activity depends on not only direct cytotoxicity but also on the modulation of the adaptive immunity through the release of immunomodulatory effectors such as IFN-γ.^49^ In this model, NaMiX promoted the generation of cytotoxic NK cells and their infiltration into BxPC-3 xenografts. However, although TriKE exhibited a small tendency to delay tumor growth by itself, highlighting a potential benefit of NK cell crosslinking to PDAC cells in vivo, NaMiX alone did not significantly delay tumor growth. This is in contrast with a previous study reporting that IL-15 therapy promotes anti-tumor immunity in a murine PDAC model ^13^ through IL-15-mediated activation of murine NK cells, limiting therefore the translational application of these results in human. Achieving optimal NK cell activation while avoiding exhaustion or tolerance, or the metabolic defects associated with continuous IL-15 treatment is critical.^50^ In this study, detailed functional and phenotypic analyses of human NK cells revealed a complex NaMiX-induced balance, between activating and inhibitory signals, that differed between splenic and intratumoral NK cell populations. In NaMiX-treated mice, splenic NK cells accumulated perforin- and granzyme-containing secretory granules, but exhibited no enhancement in degranulation or cytotoxicity against BxPC-3 cells. Phenotypically, these cells displayed increased expression of key inhibitory receptors such as NKG2A, inhibitory KIRs and TIM-3, suggesting that repeated NaMiX administrations might promote a tolerant state in peripheral NK cells. In the tumor however, infiltrated NK cells displayed an anti-tumoral NK cell phenotype, notably through the upregulation of NKG2C, an activating receptor able to stimulate strong NK cell activation and whose expression is associated with adaptive NK cells following human cytomegalovirus (CMV) exposition,^51^ and the downregulation of KLRG1, an inhibitory receptor associated with NK cell exhaustion in colorectal cancer.^52^ Altogether, these results suggest that NaMiX-stimulated infiltrated NK cells have preserved anti-tumor effector’s properties, and that a more localized administration of the treatment at the tumor site might allow to decrease peripheral inhibition while improving the functions of intra-tumoral NK cells. This humanized model was nevertheless not used for a proof of concept efficacy study, since it is limited by the low percentage of circulating NK cells.

Although NaMiX increased by multiple folds the percentage of tumor-infiltrating NK cells, they still represented only 2.4% of total xenograft cells, which limits the ability to demonstrate a therapeutic anti-tumor effect. Indeed, our in vitro data suggest that TriKE is particularly efficient against spheroids and PDOs when NaMiX-activated NK cells are readily available at the tumor site and in a large quantity. Nonetheless, this humanized model is reflective of the clinical setting, as low intra-tumoral NK cell infiltration is a hallmark of PDAC.^6^ Studies have reported that the level of peripheral and tumor-infiltrating NK cells correlates with positive clinical outcomes in PDAC.^53, 54^ Importantly, allogenic NK cell administrations in PDAC patients have resulted in promising outcomes in clinical trials.^55^ Therefore, adoptive NK cells transfer would represent an interesting approach to combine with NaMiX and TriKE stimulation, using therapeutic modalities that still need to be defined, in order to increase the number of cytotoxic tumor infiltrating NK cells. The use of chemokine^56^ or the dipeptidyl peptidase (DPP) inhibitor BXCL701^57^ could also represent a strategy to improve the efficacy of our strategy to enhance NK cell infiltration. Moreover, while an increased number of tumor-infiltrating NK cells was observed upon treatment with NaMiX, whether these NK cells are able to freely navigate or whether they are sequestered within specific regions of the xenograft remains unknown. It has recently been shown that NK cells are co-localized with tumor cells in epithelial-ductal regions in PDAC.^58^ Neutralization of ECM-NK cell interactions with antibodies against CD44, expressed in NK cells, mitigated NK cell binding to ECM components and increased NK cell invasion.^59^ The development of patient-derived orthotopic in vivo models would allow to capture NK cell localization within the TME upon treatment with NaMiX and TriKE and to characterize more precisely the complex inhibitory signals provided by the immunosuppressive cells of the PDAC TME (such as cancer-associated fibroblasts, myeloid-derived suppressor cells, tumor-associated macrophages…). ^3^ In conclusion, our results open new avenues for optimizing NK-cell-based therapeutic strategies against PDAC.

## ETHICS APPROVAL AND CONSENT TO PARTICIPATE

The study was approved by the LIH Institutional Review Board (i2TRON DTU PRIDE), and conducted in accordance with the Declaration of Helsinki. Peripheral blood mononuclear cells (PBMCs) were provided by healthy donors of the Luxembourg Red Cross giving their PBMCs for research in an anonymized way without the requirement of written informed consent. The study was approved by the Luxembourg Red Cross under the project’s number approval LIH-2018-0005. PDAC organoids were established from a tumor specimen of a patient followed at Strasbourg University Hospital under the ethics approval number CE-2022-49. The participant was informed and did not object to participation in the research or to the use of their data. The study was registered in the public project repository of the Health Data Hub (N° F20220413120650). The animal protocol was evaluated and approved by the Luxembourg Ministry of Agriculture and the Luxembourg Ministry of Health (protocol LUPA 2024/08) following the guidelines of the Directive 2010/63/EU of the European Parliament and of the Council on the protection of animals used for scientific purposes.

## CONSENT FOR BIORXIV PUBLICATION

All authors have seen and approved the manuscript, The manuscript has not been accepted or published elsewhere.

## AVAILABILITY OF DATA AND MATERIAL

Data are available upon reasonable request.

## AUTHOR’S CONTRIBUTIONS

Conception and design: CR, JZ, CSD; Methodology: CR, AP, GI, AO, J-YS, VB, CH; Analysis of data: CR, AP, GI, JYS, CSD; Writing, review and editing of the manuscript: CR, AP, GI, AO, J-YS, VE-K, VB, CH, AG, Y-JK, JZ, CSD.

## ACKNOWLEDGEMENTS

We thank the Red Cross of Luxembourg for providing access to buffy coats from healthy donors as well as the healthy donors for their blood donation. We also thank the PDAC patient for providing access to PDAC tumor cells in order to generate the organoids. The in vivo experimental timelines were created in BioRender.com.

## SUPPLEMENTAL METHODS

### Antibodies

The list of antibodies used for ELISA and flow cytometry experiments is provided below.

| Antibody | Provider | Identifier<br>(Catalog number; RRID) |
| --- | --- | --- |
| 6-His Tag Polyclonal Antibody | Bethyl Laboratories | A190-114A; AB_67321 |
| Alexa Fluor 488 Mouse anti-Ki-67 | BD Biosciences | 558616; AB_647087 |
| AlexaFluor488 Anti-6x His Tag antibody | Abcam | Ab1206; AB_298815 |
| Anti-human CD314 (NKG2D) Antibody | Biolegend | 320802; AB_492956 |
| Anti-human CD335 (NKp46) Antibody | Biolegend | 331902; AB_1027637 |
| Anti-human CD337 (NKp30) Antibody | Biolegend | 325202; AB_756106 |
| Anti-polyHistidine–Peroxidase Antibody | Sigma-Aldrich | A7058; AB_258326 |
| APC anti-His Tag Antibody | Biolegend | 362605; AB_2715818 |
| APC anti-human/mouse Granzyme B | Biolegend | 372204; AB_2687028 |
| BUV395 Mouse Anti-Human CD4 | BD Biosciences | 564724; AB_2738917 |
| BUV496 Mouse Anti-Human CD3 | BD Biosciences | 612940; AB_2870222 |
| BUV737 Mouse Anti-Human CD16 | BD Biosciences | 612786; AB_2833077 |
| BV421 anti-prpS6 phospho (Ser235/Ser263) | Biolegend | 608609; AB_2814450 |
| BV421 Mouse Anti-Human CD107A | BD Biosciences | 562623; AB_2737685 |
| BV421 Mouse Anti-Human CD45 | BD Biosciences | 563879; AB_2744402 |
| BV421 Mouse Anti-Stat5 (pY694) | BD Biosciences | 562077; AB_10894188 |
| BV605 anti-human CD11c | Biolegend | 301635; AB_2562191 |
| BV605 Mouse Anti-Human CD337 (NKp30) | BD Biosciences | 563384; AB_2738170 |
| BV711 Mouse Anti-Human CD8 | BD Biosciences | 563677; AB_2744463 |
| BV786 Mouse Anti-Human CD56 | BD Biosciences | 564058; AB_2738569 |
| FITC Mouse Anti-Human IFN-γ | BD Biosciences | 552887; AB_394516 |
| FITC Rat Anti-Mouse CD45 | BD Biosciences | 553080; AB_394610 |
| Goat Anti-Human IgG Fc (HRP) | Abcam | ab97225; AB_10680850 |
| IL-15 Monoclonal Antibody | eBioscience | 16-0157-82; AB_10596500 |
| PE anti-human CD19 Antibody | Biolegend | 302208; AB_314238 |
| PE-Cy5 CD14 Monoclonal Antibody | eBioscience | 15-0149-42; AB_2573058 |
| PE-Cy5 CD19 Monoclonal Antibody | eBioscience | 15-0199-42; AB_10853658 |
| PE-Cy7 anti-human CD11b | Biolegend | 301321; AB_830643 |
| PE-Cy7 anti-human CD314 (NKG2D) Antibody | Biolegend | 320812; AB_AB_2234394 |
| PerCP-Cy5.5 anti-human CD66b | Biolegend | 305107; AB_2077856 |
| PerCP-Cy5.5 Mouse Anti-Human Perforin | BD Biosciences | 563762; AB_2738409 |
| StrepMAB-Classic | Iba | 2-1507-001; AB_513133 |
| BUV395 Mouse Anti-Human CD183 (CXCR3) | BD Biosciences | 565223; AB_2687488 |
| BV510 anti-human CD69 Antibody | Biolegend | 310936; AB_2563834 |
| BV510 anti-human CD244 (2B4) Antibody | Biolegend | 329534; AB_2814196 |
| BV510 anti-human CD279 (PD-1) Antibody | Biolegend | 329932; AB_2562256 |
| BV605 anti-human CD62L Antibody | Biolegend | 304834; AB_2562130 |
| BV605 anti-human KLRG1 (MAFA) Antibody | Biolegend | 368626; AB_3662365 |
| BV650 anti-human CD38 Antibody | Biolegend | 303562; AB_3662213 |
| BV650 anti-human CD158e1 (KIR3DL1) Antibody | Biolegend | 312730; AB_3083305 |
| BV650 anti-human CD366 (Tim-3) Antibody | Biolegend | 345028; AB_2565829 |
| PerCP-Cy5.5 CD158a/h/g Monoclonal Antibody | eBioscience | 45-1589-42; AB_1311133 |
| PerCP-Cy5.5 anti-human TIGIT Antibody | Biolegend | 372718; AB_2632933 |
| PE anti-human CD159c (NKG2C) Antibody | Biolegend | 375004; AB_2888871 |
| PE anti-human CD57 Antibody | Biolegend | 359612; AB_2562759 |
| PE anti-human CD45 Recombinant Antibody | Biolegend | 384408; AB_3083384 |
| PE-CF594 Mouse Anti-Human HLA-DR | BD Biosciences | 568651; AB_3684436 |
| PE/Dazzle_594 anti-human CD159a Antibody | Biolegend | 375122; AB_2888869 |
| PE-CF594 Mouse Anti-Human TACTILE (CD96) | BD Biosciences | 570070; AB_3685499 |
| PE-Cy7 anti-human CD226 (DNAM-1) Antibody | Biolegend | 338316; AB_2616645 |
| PE-Cy7 anti-human CD182 (CXCR2) Antibody | Biolegend | 320716; AB_2564597 |
| APC anti-human CD328 (Siglec-7) Antibody | Biolegend | 347706; AB_2800927 |

### PDAC Organoid media composition

Basal medium:

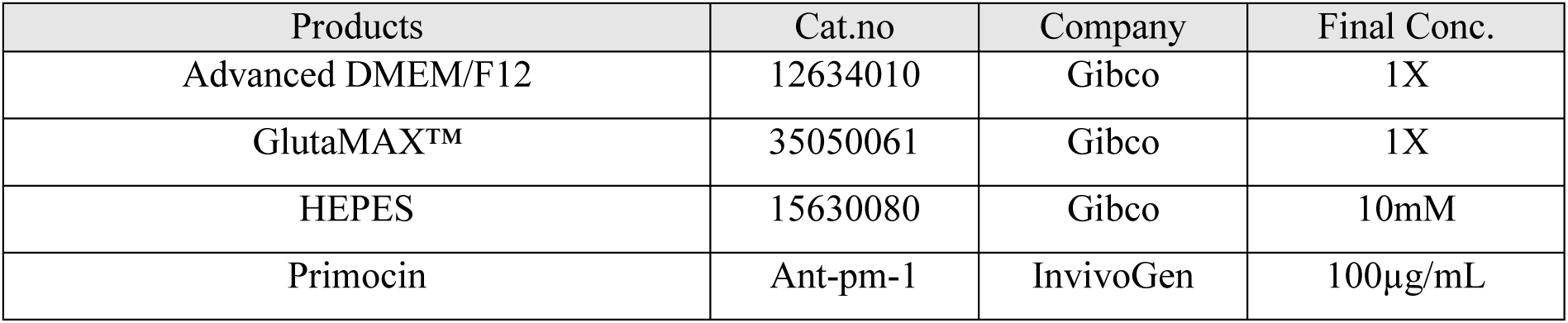

PDAC Organoid media composition Basal medium:
| Products | Cat.no | Company | Final Conc. |
| --- | --- | --- | --- |
| Advanced DMEM/F12 | 12634010 | Gibco | 1X |
| GlutaMAX™ | 35050061 | Gibco | 1X |
| HEPES | 15630080 | Gibco | 10mM |
| Primocin | Ant-pm-1 | InvivoGen | 100µg/mL |

PDAC organoid complete medium:

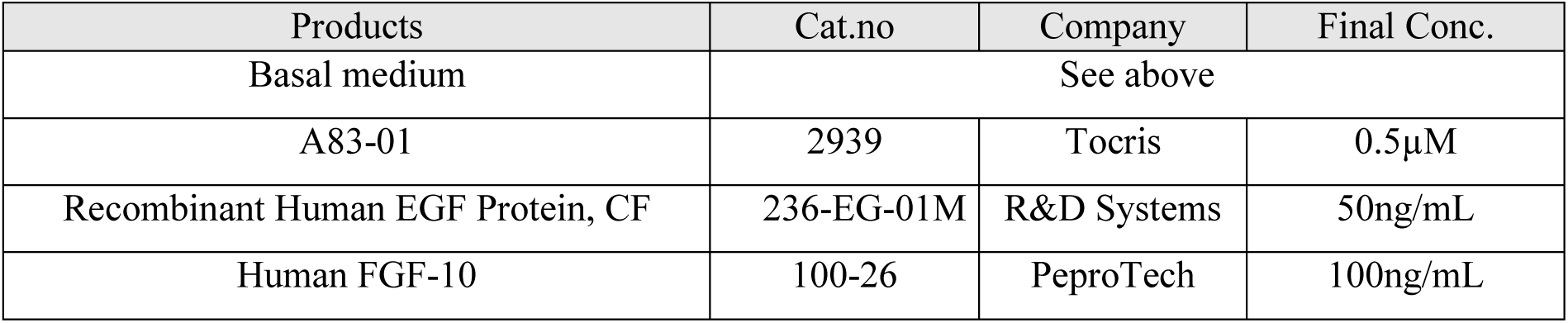
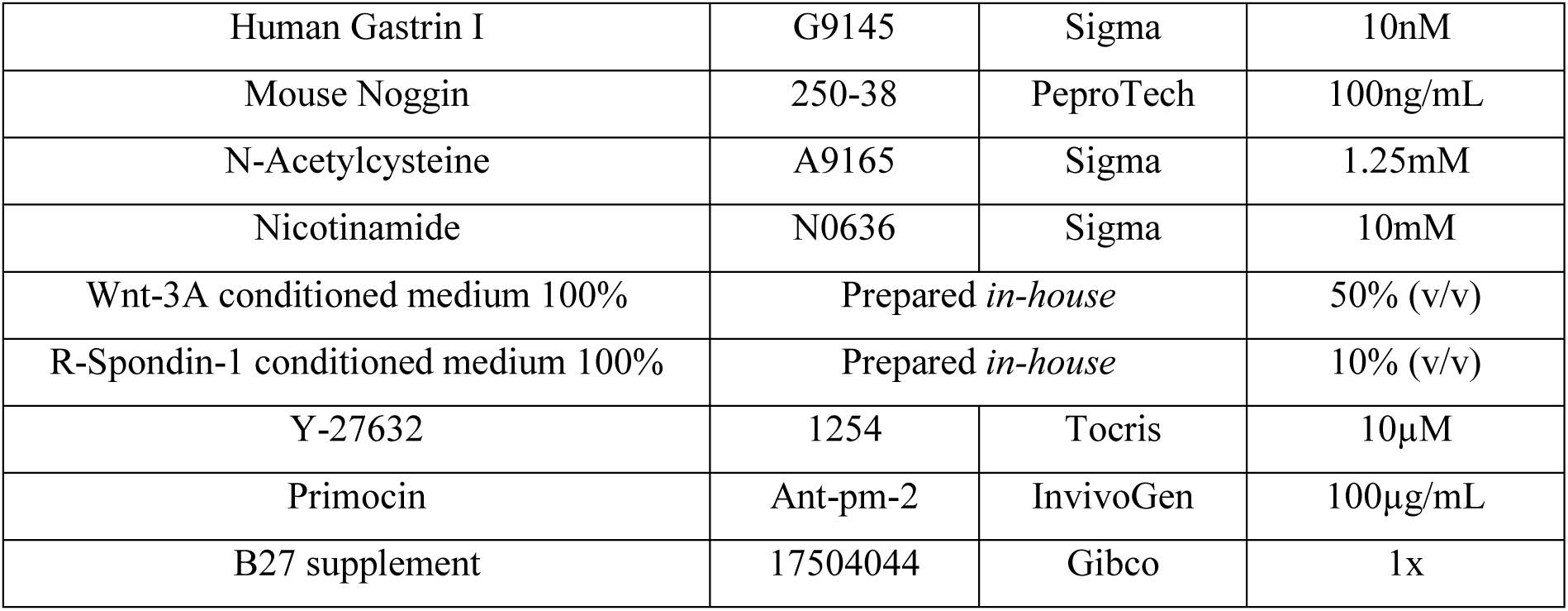

| Products | Cat.no | Company | Final Conc. |
| --- | --- | --- | --- |
| Basal medium | See above |  |  |
| A83-01 | 2939 | Tocris | 0.5µM |
| Recombinant Human EGF Protein, CF | 236-EG-01M | R&D Systems | 50ng/mL |
| Human FGF-10 | 100-26 | PeproTech | 100ng/mL |
| Human Gastrin I | G9145 | Sigma | 10nM |
| Mouse Noggin | 250-38 | PeptoTech | 100ng/mL |
| N-Acetylcysteine | A9165 | Sigma | 1.25mM |
| Nicotinamide | N0636 | Sigma | 10mM |
| Wnt-3A conditioned medium 100% | Prepared *in-house* |  | 50% (v/v) |
| R-Spondin-1 conditioned medium 100% | Prepared *in-house* |  | 10% (v/v) |
| Y-27632 | 1254 | Tocris | 10μM |
| Primocin | Ant-pm-2 | InvivoGen | 100μg/mL |
| B27 supplement | 17504044 | Gibco | 1x |

### Molecular design of constructs

The following cDNA constructs were codon optimized and synthesized (ProteoGenix SAS, Schiltigheim) for expression of the different molecules. **NaMiX**: human (hu) IL-15Rα-Sushi (UniProt n°Q13261, aa 31–205)-hu C4bp C-terminal β chain (UniProt n°P20851, aa 137–252)-scFv human scFv anti-Nkp46-6x His; **TriKE**: human scFv anti-NKG2D (patent n° US20110150870A1)-hu C4bp C-terminal β chain -human VHH anti-CEA (patent n°US2016/0083476A1)-6x His tag + human scFv anti-NKp30 (patent n°WO2021/143858A1))-hu C4bp C-terminal β chain -human VHH anti-CEA (patent n°US2016/0083476A1)-StrepXT tag; **BiKE NKG2D**: human scFv anti-NKG2D-hu C4bp C-terminal β chain-human VHH anti-CEA-6x His tag; **BiKE NKp30**: human scFv anti-NKp30 hu C4bp C-terminal β chain -human VHH anti-CEA -6x His tag.

### Establishment of cell lines and molecules production

The establishment of stable cell lines for molecules production were obtained as previously described with a few modifications.^27^ Briefly, for NaMiX, HEK293F cells were co-transfected with jetPRIME transfection reagent (Polyplus Sartorius, # 101000001) with the bi-cistronic pEFIRESpac vector coding for (hu) IL-15Rα-Sushi- hu C4bp C-terminal β chain - human scFv anti-NKp46 - 6x His and the pcDNA3.1 coding for rhu-IL15. For the TriKE and control BiKEs, HEK293T cells were (co-)transfected with the bi-cistronic pEFIRESpac vectors coding for the sequences detailed above. 48h after transfection, cells were transferred in a selection medium with the appropriate antibiotics ranging from 5–20 μg/ml of puromycin (InvivoGen, # ant-pr-1) and 100–500 μg/ml geneticine disulfate (G418) (Carl Roth, # 2039.2). Supernatants from clone culture were screened by ELISA using anti-IL-15/anti-His sandwich ELISA for NaMiX molecules and anti-StrepTag/anti-His, recombinant NKG2D (rNKG2D)/anti-His or recombinant NKp30 (rNKp30)/anti-His sandwich ELISA for TriKE, BiKE NKG2D and BiKE NKp30, respectively. The clones expressing the highest levels of molecules were expanded and transferred in suspension culture in Expi293 medium (Gibco, #A14351-01) at a density of 400.000 cells/mL. When cells reached a density of 5-6 million cells/mL, the cultured supernatant was collected, cleared by centrifugation, and filtered using 0.45 μm PVDF 1L vacuum filter units (VWR, #514-1051). Imidazole (Sigma-Aldrich, # I2399-100G) was added to culture supernatant to reach 20 mL final concentration which was then loaded on a Nickel His-Trap Excel column (Cytiva, # 29048586) over 48 h on a peristaltic pump at a flow rate of 1 ml/min and eluted on NGC chromatography system (Biorad). Purified molecules were washed and concentrated on Amicon Ultra Centrifugal Filter 50 kDa MWCO (Millipore, # UFC905008). For the TriKE, a second round of purification was performed on the His-purified solution using StrepTrapXT columns (Cytiva, #29401317).

### Western Blot

Culture supernatant of cells producing NaMiX, TriKE and BiKEs were loaded onto 4-15 % Mini-PROTEAN Tris-Glycine Extended (TGX) precast gels (Bio-Rad, # 4561086) under non-reducing conditions in Laemmli Sample Buffer (Biorad,# 1610737EDU) or under reducing conditions (using 10 % β-mercaptoethanol). Proteins were separated by electrophoresis and transferred to a polyvinylidene difluoride (PVDF) membrane by wet transfer. The membrane was blocked with 5% BSA in PBS overnight and incubated for 1 hour with AlexaFluor488 anti-His antibody (Invitrogen, #A28175). Protein bands were detected using an Amersham Typhoon 200 biomolecular image (Cytiva).

### Binding assay by ELISA

For binding assays, 1 μg/mL of the appropriate recombinant (r) protein rNKp46 (R&D Systems, #1850-NK-025), rNKp30 (Sinobiological, #10480-H02H), rNKG2D (Sinobiological, #10575-H01S) or rCEACAM5 (rCEA) (R&D systems, #10449-CM) was coated on a MaxiSorp 96-well flat-bottom ELISA plate (ThermoScientific, # 442404) overnight. All incubations with antibodies were done for 1 hour at 4℃, washed using 1% PBS/BSA (Carl Roth, #1ET9.1) and blocked with 5% PBS/BSA. Decreasing construct (NaMiX, TriKE or BiKE) concentrations ranging from 20 ug/mL to 0 ug/mL were added (in absence or in presence of 20 μg/mL of blocking antibodies) and the revelation was performed using Rabbit anti-His-peroxidase antibody (Sigma-Aldrich, # A7058-1VL).

### Fluorescence and confocal microscopy

KHYG-1 cells and BxPC-3 cells were stained with CellTracker Deep Red (Invitrogen, #C34565) and CellTrace CFSE (Invitrogen, #C34554) respectively, following manufacturer’s instructions. Both cell types (60,000 cells each/condition) were co-incubated with 5 μg of TriKE, previously stained with AlexaFluor 594 Microscale Protein Labeling Kit (Invitrogen, #A30008) for 2 hours at 37°C. 30 minutes before the end of the incubation, Hoechst 33342 (Miltenyi Biotec, #130-111-569) was added to all conditions. Cells were then transferred to poly-L-lysine-coated µ-slide chambers (Ibidi #80806) and fixed with 2% paraformaldehyde for 20 minutes at 4°C. Cells were gently washed with PBS, and the medium was replaced with Ibidi mounting medium (Ibidi, #50001). Images were acquired using a Zeiss Observer Z1 fluorescence microscope or a Zeiss LSM880 fast Airyscan confocal microscope with excitation lasers 488, 594, and 633 nm.

### Humanized mice studies

NOD.Cg-Prkdcscid Il2rgtm1Wjl/SzJ (NSG) mice, aged 4 week, were first humanized. 48h before stem cell engraftment, the mice were injected with Busulfan (1,4-Butanediol dimethanesulfonate) (Merck) diluted in PBS (1,2 mg/mL) intraperitoneally at a dose of 20 mg/kg in a total volume of 10 mL/kg. 24h before stem cell engraftment, a second dose of Busulfan (identical to the first one) was given to the mice intraperitoneally. On the day of engraftment, NSG mice were injected intravenously in the tail vein with 50,000 human CD34^+^ hematopoietic stem cells (HSC) derived from umbilical cord (Lonza, Belgium) in FBS-free RPMI medium. Around 12 weeks post CD34^+^ cells administration), mice were injected with 2,5 μg human recombinant IL-15 (Peprotech, #200-15-10UG) + 7,5 μg of human recombinant IL15Rα (Peprotech, #200-15RA-100UG) intraperitoneally. Humanization was achieved 20 weeks post CD34^+^ cells administration (humanization rate > 40%). BxPC-3 cells were then harvested during exponential growth and injected subcutaneously in the right flank of mice (2 million cells in 100 μl FBS-free RPMI/mouse). Two weeks post-transplantation, the mice were randomized based on the tumor size and humanization rate and allocated into the experimental groups with 5-7 mice in each group. Depending on the group, mice received intraperitoneal injections of the molecules at a dose of 1 mg/kg (vehicle group received PBS). Tumor growth was monitored twice weekly by Vernier digital caliper. The tumor volume was calculated as followed: Volume = 0.52 x l x w^2^ (l: length of the longest diameter (mm), w=length of the axis perpendicular to l (mm)). At the end of the experiment, mice were anaesthetized and euthanized by intracardiac terminal blood puncture followed by a PBS flush. The spleen was removed and cells were isolated. Tumors were excised and dissociated with the Tumor Dissociation Kit (Miltenyi Biotec, # 130-095-929) following manufacturer’s instructions to retrieve cells. All cell suspensions were then blocked using Human TruStain FcX Fc Receptor Blocking solution (Biolegend, # 422301) for 5 minutes at 4°C and then stained with a Live/Dead and with antibody panel containing CD3, CD4, CD8, CD16, CD56, CD14, CD19, CD11c, CD66b and CD11b for study of immune cell subpopulations. For the ex vivo experiment, 500.000 splenic cells were incubated at a ratio 10:1 with CellTrace Violet-stained BxPC-3 cells for 5 hours following the protocol in the “NK cell cytotoxicity assays by flow cytometry” paragraph of the main material and methods section. Acquisition was performed on the LSR Fortessa flow cytometer (BD Biosciences) and analyzed with Kaluza (Beckman Coulter). t-SNE analysis were performed on CellEngine (cellengine.com).

## SUPPLEMENTAL FIGURES

**Supplemental Figure 1:**
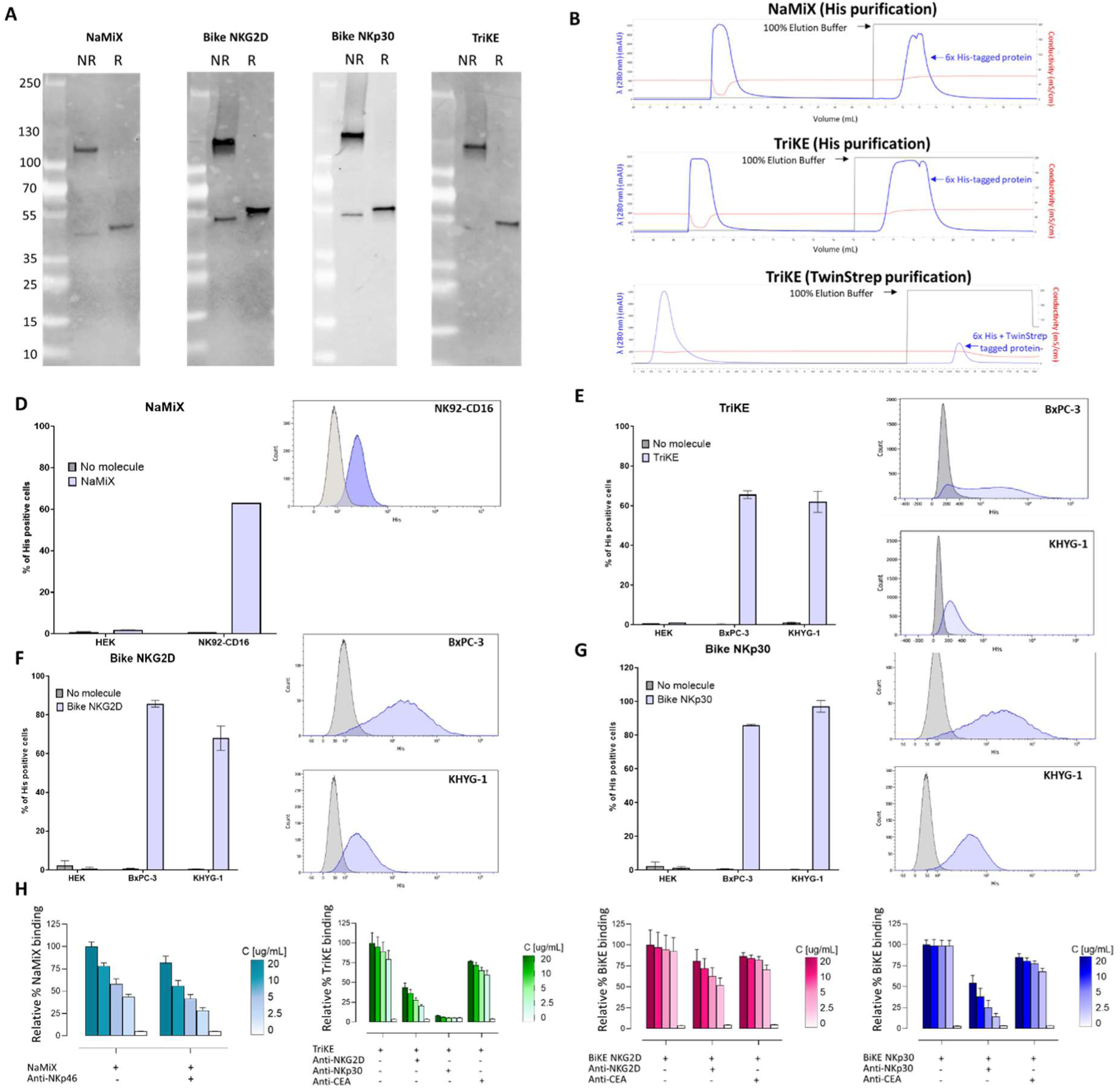
Molecular characterization and purification of NaMiX and Engagers. (A) The molecular pattern of NaMiX, TriKE and control BiKEs was analyzed by Western Blot under non-reducing (NR) or reducing (R) conditions and revealed with anti-His antibody coupled to AlexaFluor 488. (B) NaMiX and TriKE were purified using His affinity chromatography and TriKE was further purified using StrepXT affinity chromatography. Binding of NaMiX (D), TriKE (E), BiKE NKG2D (F) and BiKE NKp30 (G) was evaluated on HEK-293T (CEA^-^NKG2D^-^NKp30^-^), BxPC-3 (CEA^+^), NK92-CD16 (NKp46^+^) and on KHYG-1 (NKG2D^+^ NKp30 ^+^) cell lines and detected by anti-His antibody by flow cytometry. (E-F) Binding of NaMiX, TriKE and BiKEs was also assessed by ELISA. (H) For NaMiX, recombinant human NKp46 (rNKp46) was coated then NaMiX was added in presence (red) or in absence (green) of blocking antibody and revelation was performed using anti-6x his antibody conjugated to HRP. For TriKE and BiKEs, ELISA assays were performing using rNKG2D, rNKp30 or rCEA as coating, then adding the engager of interest in presence or absence of adequate blocking antibody (anti-NKG2D, anti-NKp30 or anti-CEA) then revealing with anti-6x his antibody conjugated to HRP. Data are presented as the mean values ± SEM. Results correspond to two pooled independent experiments (2-3 replicates per experiment).

**Supplemental Figure 2:**
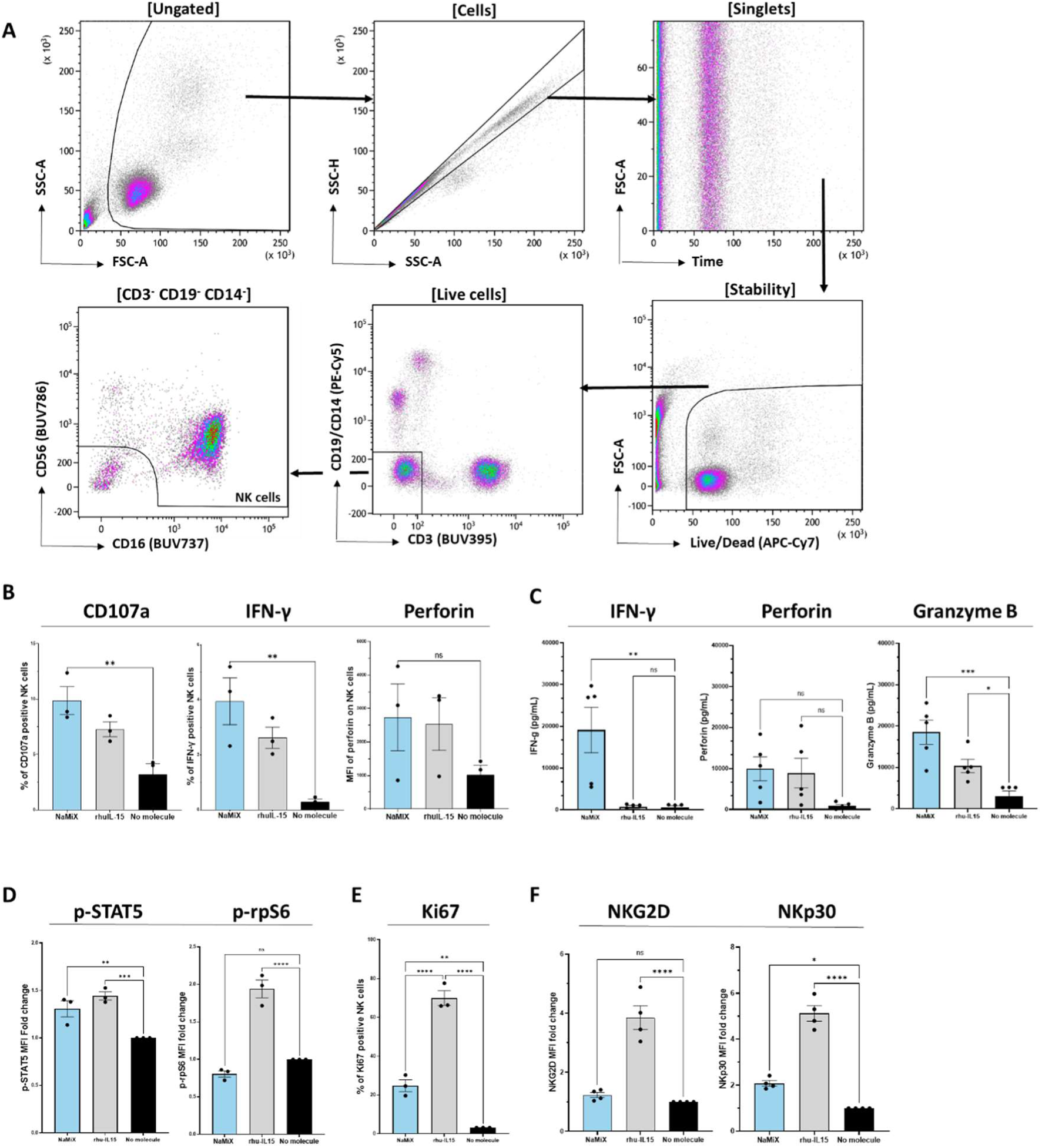
NaMiX stimulates the degranulation and activation of NK cells via STAT5 but not mTOR pathway. Healthy donors PBMCs were pre-incubated with NaMiX, rhu-IL15 or control medium for 48 hours. PBMCs were stained with LIVE/DEAD™ Fixable Near-IR Dead Cell Stain Kit (to exclude dead cells), anti-human CD3 (to exclude CD3^+^ T cells), anti-human CD14 and CD19 (to exclude B cells and monocytes), and with anti-human CD56 and CD16 to identify NK cells. (A) Gating strategy. (B) NK cells were then analyzed by flow cytometry for their expression of CD107a, IFN-γ and perforin. (C) The cell culture supernatant was analyzed for perforin, granzyme B and IFN-γ by ELISA. (D) PBMCs were incubated with NaMiX for 5 minutes then NK cells were analyzed by flow cytometry for their expression of p-STAT5 and p-rpS6. (E-F) PBMCs pre-incubated with NaMiX for 48 hours were studied for their expression of Ki67, NKG2D and NKp30 on NK cells by flow cytometry. Data are presented as the mean values ± SEM. Results correspond to two pooled independent experiments (1-3 donors per experiment). Statistical analysis was performed using a one-way ANOVA and post-hoc Tukey test (*p<0.05; **p<0.01; ***p<0.001; ****p<0.0001).

**Supplemental Figure 3:**
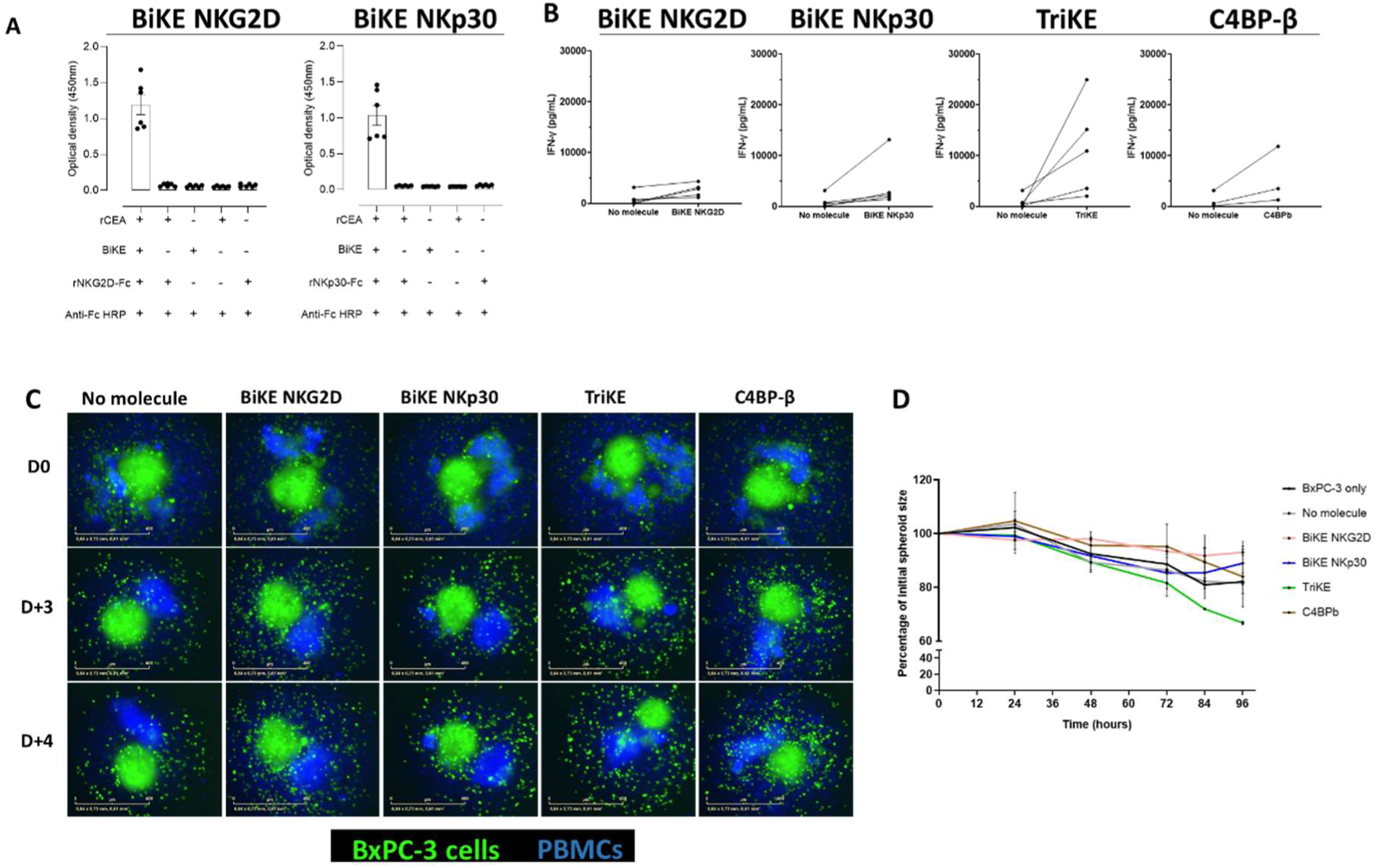
Crosslinking and NK cell activation by BiKEs and TriKE. (A-B) ELISA assay using recombinant human CEA (rCEA) as coating, adding BiKE, then revealing by (A) recombinant NKG2D coupled to Fc (rNKG2D-Fc) or (B) recombinant NKp30 coupled to Fc (rNKp30-Fc) and an anti-Fc linked with HRP. Data are presented as the mean values ± SEM. Results correspond to two pooled independent experiments (3 replicates per experiment). (B) PBMCs were incubated with BiKE, TriKE or control scaffold C4BPβ together with BxPC-3 cells at an E:T ratio of 10:1 for 24 hours and the cell culture supernatant was analyzed for IFN-γ by ELISA. Data is represented by individual donor (n=3-5). (C-D) CFSE-stained BxPC-3 cells were seeded in ultra-low attachment plates for 48 hours, then co-incubated with CellTracker Deep Red stained PBMCs (blue) and BiKEs, TriKE, control scaffold C4BPβ or control medium and placed inside IncucyteS3 for 96 hours. Pictures were acquired every 6 hours and spheroid size was quantified using ImageJ software. Scale bar = 400 µm.

**Supplemental Figure 4:**
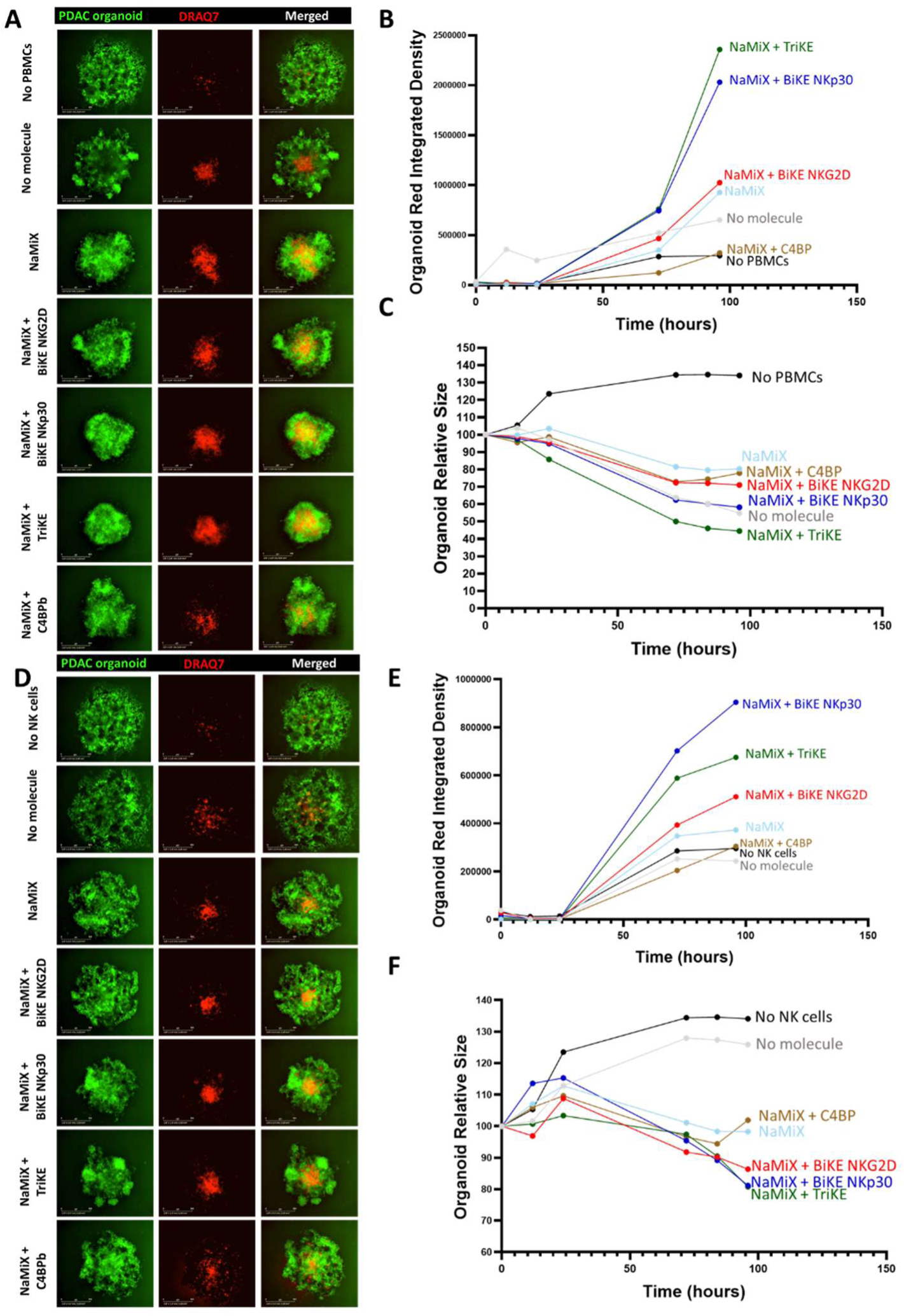
Effect of NaMiX and Engagers on PBMCs and NK cell cytotoxicity against patient-derived PDAC organoids. CFSE-stained patient-derived organoids were seeded in ultra-low attachment plates together with (top panel) PBMCs at an E:T ratio 10:1 or (bottom panel) purified NK cells at an E:T ratio 1:1 and the cytotoxicity marker DRAQ7 with indicated molecules and placed inside IncucyteS3 for 96 hours. (A, D) Representative images at day 4 of co-incubation. Incucyte 2025B software was used to quantify (B, E) the red integrated density (C, F) and organoid relative size. The experiment was performed 3 times and one representative donor is shown. Scale bar = 800 µm.

**Supplemental Figure 5:**
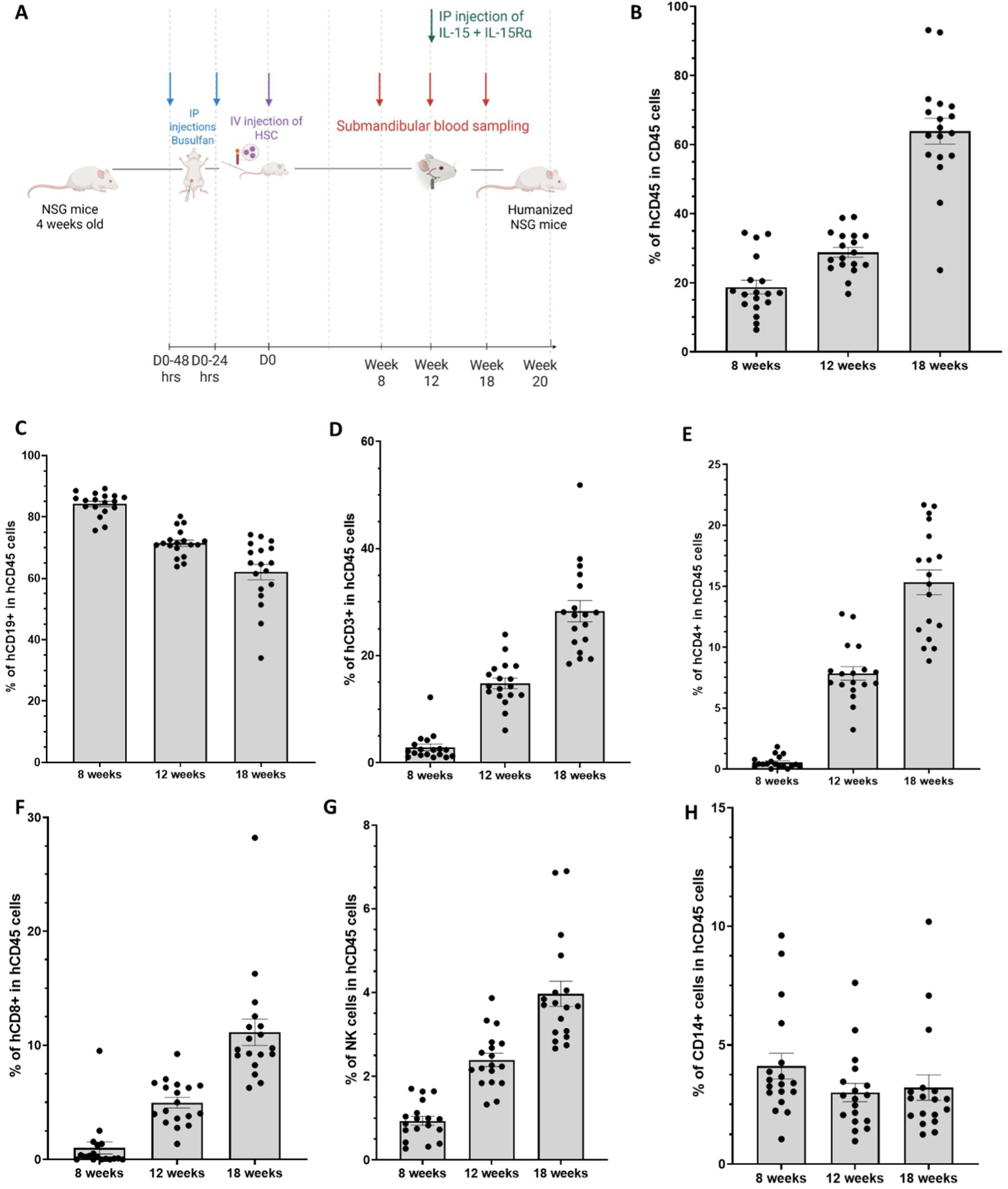
Development of the humanized mice model. (A) Schematic representation of the humanized mice model. Blood was drawn at 8, 12 and 18 weeks post-CD34^+^ hematopoietic stem cells (HSCs) engraftment and the percentages of (B) hCD45, (C) hCD19^+^ B cells, (D) hCD3^+^ T cells, hCD4^+^ T cells, (F) hCD8^+^ T cells, (G) NK cells, and (H) hCD14^+^ monocytes were quantified by flow cytometry and are expressed as percentage of hCD45. Data are presented as the mean values ± SEM. IP: intraperitoneal; IV: intravenous.

